# High-resolution comparative single-cell transcriptomics of *doublesex*-expressing neurons reveals evolutionary conservation and diversity of sexual circuits in *Drosophila*

**DOI:** 10.1101/2025.05.06.652433

**Authors:** Justin Walsh, Ian P. Junker, Yu-Chieh David Chen, Yen-Chung Chen, Helena Gifford, Yun Ding

## Abstract

Understanding how the cellular and molecular composition of neural circuits change during evolution is essential for deciphering how behavior evolves. Male sexual behaviors in *Drosophila* species are remarkably diverse, and the underlying sexual circuits are specified by sex determination genes. Here, we employed single-cell transcriptomics to systematically characterize and compare neuron cell types that express the sex determination gene *doublesex* in adult males across *Drosophila* species with divergent sexual behaviors. High-resolution profiling to delineate cellular diversity revealed a largely conserved set of cell types across four species, with minimal evolutionary gain or loss and potentially more prevalent changes in cell type abundance. In-depth comparisons between *D. melanogaster* and *D. yakuba* showed that transcriptomic conservation and differentially expressed genes between species are highly cell-type-specific, suggesting that cell types evolve as highly independent units. We identified widespread species differences in gene expression, particularly in the neuromodulatory signaling pathway, while preserving a conserved circuit layout in sexual identity and neurotransmitter properties. We further generated a female dataset in *D. melanogaster* to define sex differences in cell types and examine species divergence in relation to sex differences. Finally, we reported marker gene combinations that uniquely define each cell type, providing a foundational resource that enables the design of cell -type-specific genetic reagents. Overall, our study provides fundamental insights into the cellular diversity of sexual circuits and how evolution shapes cell types and gene expression in behavioral adaptations.

## Introduction

Even among closely related species, behaviors may vary drastically, and this behavioral variation largely arises from evolutionary changes in the structure and function of the underlying homologous neural circuits (Katz & Harris-Warrick, 1999; Katz, 2011; Roberts et al., 2022). These changes can occur in the presence and absence of specific cell types, the abundance of a particular cell type, or gene expression that influences neuron anatomy and physiology. Facilitated by the unprecedented capacity of single-cell RNA sequencing (scRNA-seq) to define molecular cell types, recent comparative studies have begun to shed light on species conservation and differences in cell types (Shami et al., 2020; Khrameeva et al., 2020; Schafer et al., 2022; Geirsdottir et al., 2019; Wei et al., 2022; Lee & Benton, 2023; Wang et al., 2024). However, existing studies often focus on distantly related species and/or have limited resolution to capture fine-scale changes in cell types. Particularly, since the basic cellular organization of the nervous system tends to be largely conserved among closely related species (Katz & Harris -Warrick, 1999, Katz, 2011), subtle changes in the nervous system may result in major behavioral differences but go undetected in existing datasets. With few exceptions of relatively simple organisms (e.g., Zhao et al., 2008; Pollo et al., 2024; Toker et al., 2024), it remains unclear how much neuronal cell types differ among closely related species with divergent behaviors at a fine-grained level. Furthermore, it is unknown whether certain cell types exhibit greater divergence across species and how gene expression evolves at the level of individual cell types.

*Drosophila* species, with their rich behavioral diversity and experimental accessibility, serve as a powerful system to study behavioral evolution through species comparisons (Prieto-Godino et al., 2017; Seeholzer et al., 2018; Ding et al., 2019; Auer et al., 2020, 2022; Takagi et al. 2024; Ye, et al., 2024; Coleman et al., 2024; Li et al., 2024). *Drosophila* sexual behaviors present a unique opportunity as the genes and neural circuits underlying these behaviors have been well studied in the model species *D. melanogaster* and these behaviors have rapidly diversified across species (Spieth et al., 1974; Yamamoto & Koganezawa, 2013; Shirangi et al., 2016; Ding et al., 2019; Shen et al., 2023; Lillvis et al., 2024). For example, *Drosophila* males perform complex and diversified courtship rituals such as singing a species -specific courtship song by vibrating wings (Ewing & Bennet-Clark 1968; Greenspan & Ferveur, 2000). Species also vary in their responses to sensory cues that are visual, auditory, and chemosensory (Ferveur, 2005; Billeter et al., 2009; Gleason et al., 2012; Seeholzer et al., 2018; Khallaf et al., 2021). For example, the sex pheromone 7-tricosene is male-specific in *D. melanogaster*, where it suppresses male courtship, but is sexually monomorphic in *D. yakuba* and instead promotes male courtship (Coleman et al., 2024).

These sexual behaviors are orchestrated by two key transcription factors in the sex determination pathway: *doublesex* (*dsx*) and *fruitless* (*fru*). The *dsx* gene is spliced into sex-specific isoforms that regulate the development and function of sexual circuits (Villella & Hall, 1996; Rideout et al., 2007; Sanders & Arbeitman, 2008; Rideout et al., 2010; Shirangi et al., 2016). Approximately 900 *dsx*+ neurons, representing about 1% of the adult male central nervous system, are distributed in anatomically and functionally distinct neuronal clusters. For instance, the male-specific TN1 cluster in the ventral nerve cord (VNC) is a group of motor patterning neurons responsible for generating courtship song (Shirangi et al., 2016); the sexually dimorphic pC1 cluster in the brain integrates sensory cues and contains functionally heterogeneous populations that encode sexual and aggressive states (Kimura et al., 2008; von Philipsborn et al., 2011; Deutsch et al., 2020; Schretter et al., 2020). Given the expanding knowledge of their roles in sexual behaviors–which evolve rapidly across species–*dsx*+ neurons provide an excellent inroad for uncovering the cellular and molecular mechanisms underlying the evolution of complex behaviors and species-specific adaptations. Indeed, we have previously linked the lineage-specific loss of sine song (a type of courtship song) in *D. yakuba* to the loss of a neuronal subtype within the TN1 cluster (Ye, et al., 2024).

In this study, we overcome the common challenge of cellular resolution in comparative scRNA-seq studies by profiling genetically labeled *dsx*+ neurons across *Drosophila* species. This effort resulted in a scRNA-seq atlas comprising 46,922 *dsx*+ neurons, with each *dsx*+ neuron represented an average of 52.1 times. The dataset included four species within the *D. melanogaster* subgroup: *D. melanogaster* (19.1x coverage), *D. yakuba* (20.1x), *D. santomea* (6.0x), and *D. teissieri* (7.1x) (**Supplemental Table 1**)—a group of closely related species that exhibit a wide range of differences in sexual behaviors and are emerging models for comparative studies of the nervous system and behavior (Stern et al., 2017; Ding et al., 2019; Coleman et al., 2024; Ye, et al., 2024; Li et al., 2024). In total, we molecularly defined 84 cell types, all systematically mapped to and hierarchically organized by known *dsx*+ neuronal clusters, providing a comprehensive characterization of cellular diversity within this neuronal population of broad interests. We found that sexual circuits evolve through widespread cell-type-specific gene expression changes on top of a largely conserved organization in cell types, sex identity, and neurotransmitter properties. We also generated a scRNA-seq dataset in adult *D. melanogaster* females to characterize female *dsx*+ cell types and explore the relationship between species and sex differences. Finally, we compiled a list of marker gene combinations that uniquely target each cell type and hosted our datasets on a Shiny app interface that allows user-defined tasks such as gene expression inquiries and molecular marker detection across cell types, species, and sexes. Altogether, our study offers both novel insights and useful resources for understanding and dissecting the function and evolution of neural circuits.

## Results

### High-resolution characterization of cell type diversity within *dsx+* neurons

We genetically labeled *dsx*+ neurons using CRISPR/Cas9-mediated genome editing (Ye et al., 2024) and fluorescently sorted them from adult male brains and VNCs of four *Drosophila* species that diverged within the past 12 million years (**Figure 1A**; Tamura et al., 2004; Bachtrog et al., 2006). In total, we generated three replicates for *D. melanogaster* and *D. yakuba,* including one previously published replicate for each species (Ye, et al., 2024). We primarily focused on the comparison of cell types and gene expression between these two species of common interests (Stern et al., 2017; Ding et al., 2019; Ye, et al., 2024; Coleman et al., 2024). We also included one sample each for *D. santomea* and *D. teissieri* to provide a general perspective of cell type conservation across species and phylogenetic context for gene expression changes we identified between *D. melanogaster* and *D. yakuba* (**Figure 1A**).

**Figure 1.**
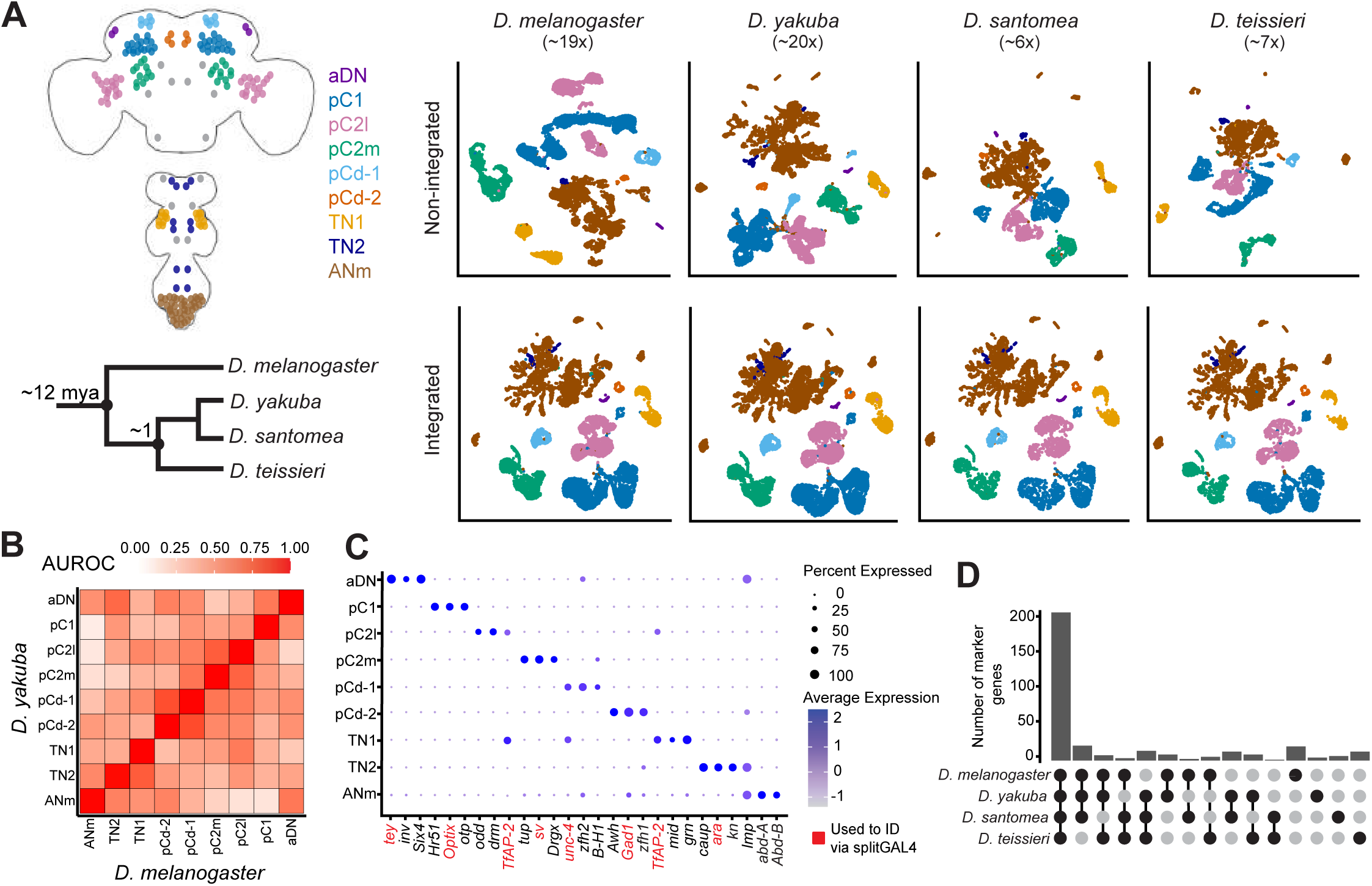
Identification of parental clusters in scRNAseq data in four species. **(A)** Left: schematic representation of the cell bodies of dsx+ neurons in the brain and VNC, color-coded by parental clusters, and a phylogeny of species included in this study (divergence times shown in millions of years ago, based on Tamura et al., 2004; Bachtrog et al., 2006). Gray dots represent dsx+ neurons that are not identified in our scRNA-seq data. Right: UMAP representations of scRNA-seq data from four species, color-coded to match the dsx+ parental clusters on the left. The top row shows single species (non-integrated) data, while the bottom row displays the integrated data separated by species. **(B)** Heatmap showing the similarity (AUROC) of parental clusters between *D. melanogaster* and *D. yakuba*. **(C)** Dotplot illustrating the average expression and percentage of cells expressing three marker genes for each parental cluster. Genes highlighted in red were used to identify parental clusters via split-GAL4. All genes shown are conserved across all four species. **(D)** Barplot displaying the number of marker genes conserved across species, including only the top 20 marker genes for each parental cluster for each species. Marker genes from all nine parental clusters are shown.

Starting with *D. melanogaster* scRNA-seq data, we performed an initial clustering with the goal of identifying previously defined *dsx*+ neuronal clusters (Lee et al., 2002; Kimura et al., 2008; Rideout et al., 2007; Sanders & Arbeitman 2008; Robinett et al., 2010; Rideout et al., 2010), which we refer to as “parental clusters” (**Figure 1A**). Upon identifying unique marker genes, we used a split-GAL4 genetic intersectional approach, pairing *dsx* hemi-drivers with hemi-drivers against marker genes for each parental cluster. This strategy, combined with previously known marker genes such as *Hox* genes, allowed us to validate the fidelity of the clustering and successfully assign parental cluster identities: aDN (*tey*+); pC1 (*Optix*+); pC2l *(TfAP-2*+, *tsh*-); pC2m (*sv*+); pCd-1 (*unc-4*+); pCd-2 (*Gad1*+); TN1 (*TfAP-2*+, *tsh*+); TN2 (*ara*+, *abd-A*-, *Abd-B*-), ANm (*abd-A*+ and/or *Abd-B*+) (**Figure 1B and 1C** and **Supplemental Figures 1 and 2A**). We did not identify cell clusters corresponding to the single-neuron pairs pMN1, pMN3, pLN, and sLG and the TN2 single-neuron pairs prA, prC, msB *(ara* partially labeled TN2), which together comprise less than 2% of all *dsx*+ neurons (cell bodies shown in gray color in **Figure 1A**). As these unidentified neurons exhibit larger soma, we speculate that they might be lost during sample preparation due to sensitivity to experimental procedures or stringency of cell sorting.

**Figure 2.**
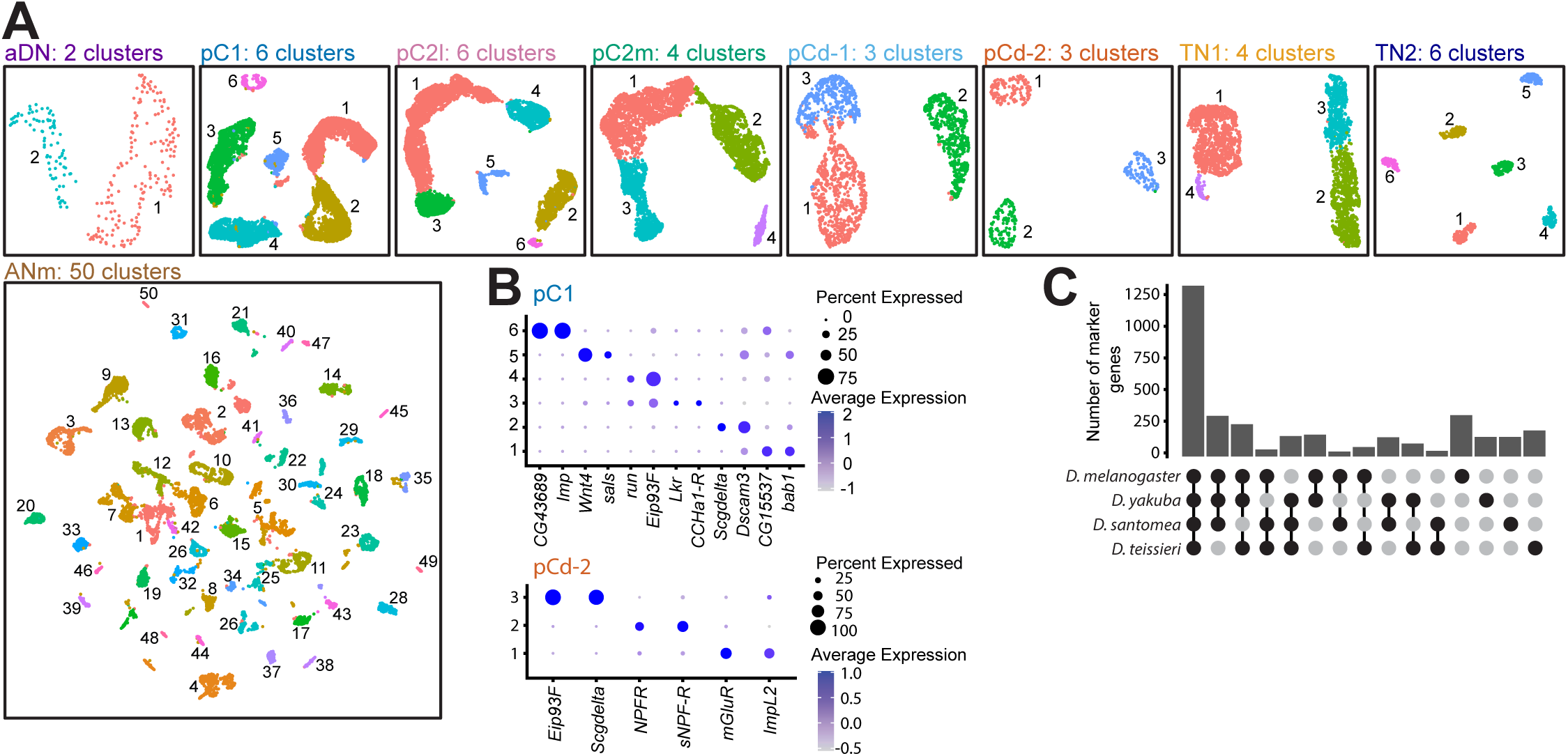
Subclustering analyses of the parental clusters from the integrated four-species dataset. **(A)** UMAP representations of the subclustering of parental clusters. Different colors represent molecularly distinct cell types. **(B)** Representative dotplots showing the average expression and percentage of cells expressing two marker genes for each subcluster within pC1 (top) and pCd-2 (bottom). **(C)** Barplot displaying the number of marker genes conserved across species, including only the top 20 marker genes for each subcluster for each species. Marker genes from all subclusters are shown.

Following this initial clustering, which faithfully recapitulated existing knowledge of *dsx*+ clusters, we transferred the identities of parental clusters in *D. melanogaster* to the other three species using the expression of the above marker genes. Furthermore, we used MetaNeighbor (Crow et al., 2018) to ensure our transferred parental identities were accurate (i.e., had the highest AUROC scores) ( **Figure 1B**, **Supplemental Figure 3**). This high degree of species consensus justified the further integration of data from all the four species into one dataset for identifying and comparing cell types on a common clustering space (**Figure 1C**). Overall, the marker genes for parental clusters were well-conserved across species. The majority of the top 20 markers for each parental cluster in a species were also marker genes for the same cluster in the other three species (**Figure 1D**).

To resolve the cellular diversity within each parental cluster, we extracted each parental cluster from the integrated dataset for subclustering and identified a total of 84 subclusters that we refer to as cell types (**Figure 2A**). The aDN, pCd-2, and TN2 clusters resolved into 2, 3, and 6 subclusters, respectively (**Figure 2A**), each likely representing a cell type corresponding to a single-neuron pair, most of which are distinguishable by a single marker gene (**Figure 2B** and **Supplemental Figure 4**). Larger parental clusters that likely originate from a single hemilineage origin, such as pC1 and pC2 (Ren et al., 2016) — whose heterogeneity has been widely appreciated but remains poorly understood—resolved into several subclusters, each typically representing a group of neurons (**Figure 2B** and **Supplemental Figure 4**). While some of these subclusters are distinguishable by a single marker (e.g., pC1_6), others (e.g., pC1_1 and pC1_3) are harder to define and often display continuity in clustering space ( **Figure 2A, 2B**). This continuity may reflect the biological nature of these cell types defined by the quantitative expression levels of multiple genes. The ANm cluster, as expected given its diverse hemilineage origins (Sanders & Arbeitman, 2008; Rideout et al., 2010), resolved into a large number (50) of more discrete subclusters, many of which are well-defined by one or two marker genes, revealing extensive cellular diversity ( **Figure 2A**). Based on described hemilineage markers (Bossing et al., 1996; Schmid et al., 1999; Lacin et al., 2019; Allen et al., 2020), we annotated the hemilineage origin for 25 of the 50 ANm subclusters, representing at least 16 distinct hemilineages (**Supplemental Figure 2**; **Supplemental Table 2**). Similar to the marker genes for parental clusters, these subcluster marker genes are largely shared across four species (**Figure 2C**). Unless stated otherwise, all subsequent analyses were conducted on these subclusters.

### Overall conservation of *dsx+* cell types across species

The systematic characterization of cell types set the stage for species comparison. Focusing on *D. melanogaster* and *D. yakuba*, where biological replicates were available, we assessed species differences in the relative abundance of cell types using scCODA, a Bayesian model designed to detect differences in cell numbers across groups based on scRNA-seq data (Büttner et al., 2021). To ensure the accuracy of our comparison, we manually quantified the number of cells in each *dsx+* parental cluster across species by immunostaining (excluding ANm due to its high density of *dsx*+ neurons, making precise counting difficult) (**Supplemental Table 3**).

We then scaled the scRNA-seq cell count for each cell type accordingly so that the size of each parental cluster aligned with the immunostaining results. Among the 84 molecularly distinct subclusters, the only species-specific one was TN1_4 (**Figure 3A** and **3B**). This subcluster corresponds to one we previously identified (Ye, et al. 2024), which is a TN1 subtype associated with the lineage-specific loss of sine song in *D. yakuba* males. We also defined 12 species-biased subclusters that varied in cell number by 1.3-to 6.8-fold. Of these, ten were in the ANm (ANm_4/8/17/18/24/25/32/33/45/48, all *D. yakuba*-biased), the other two were TN1_2 (*D. melanogaster*-biased) and pC1_6 (*D. yakuba*-biased) (**Figure 3A** and **3B**). Each of the three ANm subclusters with annotated hemilineage origins (ANm-25/32/45) originates from a distinct hemlineage (**Supplemental Figure 2B**). These subclusters are potential candidates of more variable cell types across species for future validation and functional investigation.

**Figure 3.**
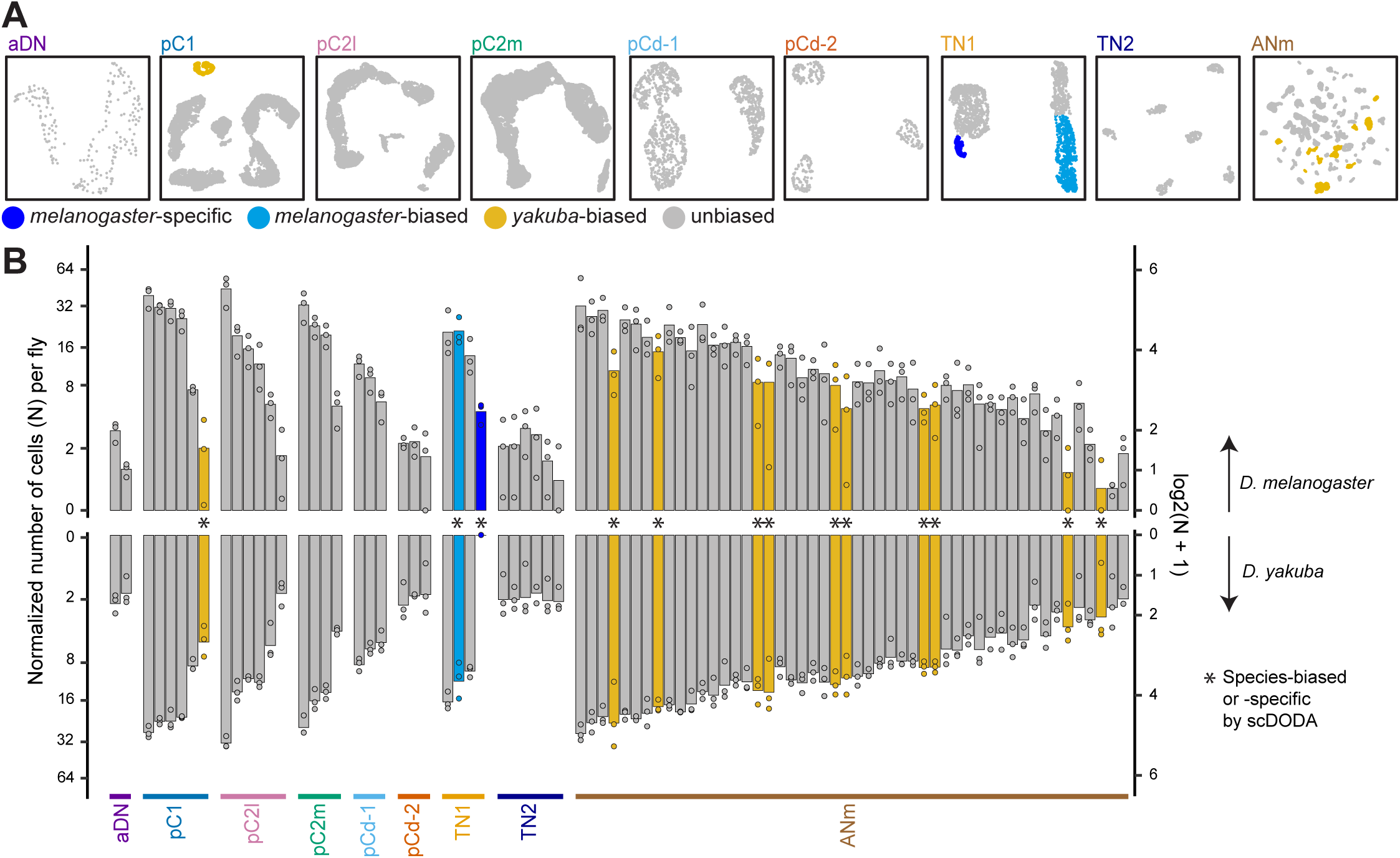
Cell type number comparison between *D. melanogaster* and *D. yakuba*. (**A**) UMAP representations of the subclustering of parental clusters, color-coded by species-biases in cell number within each subcluster. (**B**) Barplot showing the normalized number of cells (N) per fly in each subcluster, separated by species and grouped by parental cluster identity shown at the bottom. Numbers were plotted on a log2 scale with an offset of 1 (log2(N+1)) along the Y-axis to accommo-date a wide range of values and account for zeros.

Of the 83 subclusters shared between *D. melanogaster* and *D. yakuba*, 82 were present in all four species. A small subcluster in the ANm (ANm_47) was not detected in *D. santomea*, which may reflect a true species difference or could be due to its lower coverage of *dsx+* neurons. We did not perform scCODA analysis for *D. santomea* and *D. teissieri* due to the lack of biological repeats. Nevertheless, the comparison across all four species strongly indicates that species differences in the presence or absence of cell types are rare.

### Cell type-specific transcriptomic conservation and divergence across species

A neuron’s morphology and function, to a great extent, are the product of its gene expression. Therefore, we asked if some cell types are more similar or different in gene expression profile across species, potentially revealing circuit nodes with more constrained or divergent functions. For each subcluster, we used Spearman rank correlation of gene expression across species to provide a continuous measure for the degree of transcriptomic conservation (Kim et al., 2018; Skinnider et al., 2019). Subclusters with higher across-species correlation scores indicate greater transcriptomic conservation, whereas those with lower scores indicate greater transcriptomic divergence. We found that the correlation scores between *D. melanogaster* and *D. yakuba*, ranging from 0.60 to 0.73, varied substantially even among cell types within the same parental cluster (**Figure 4A**).

**Figure 4.**
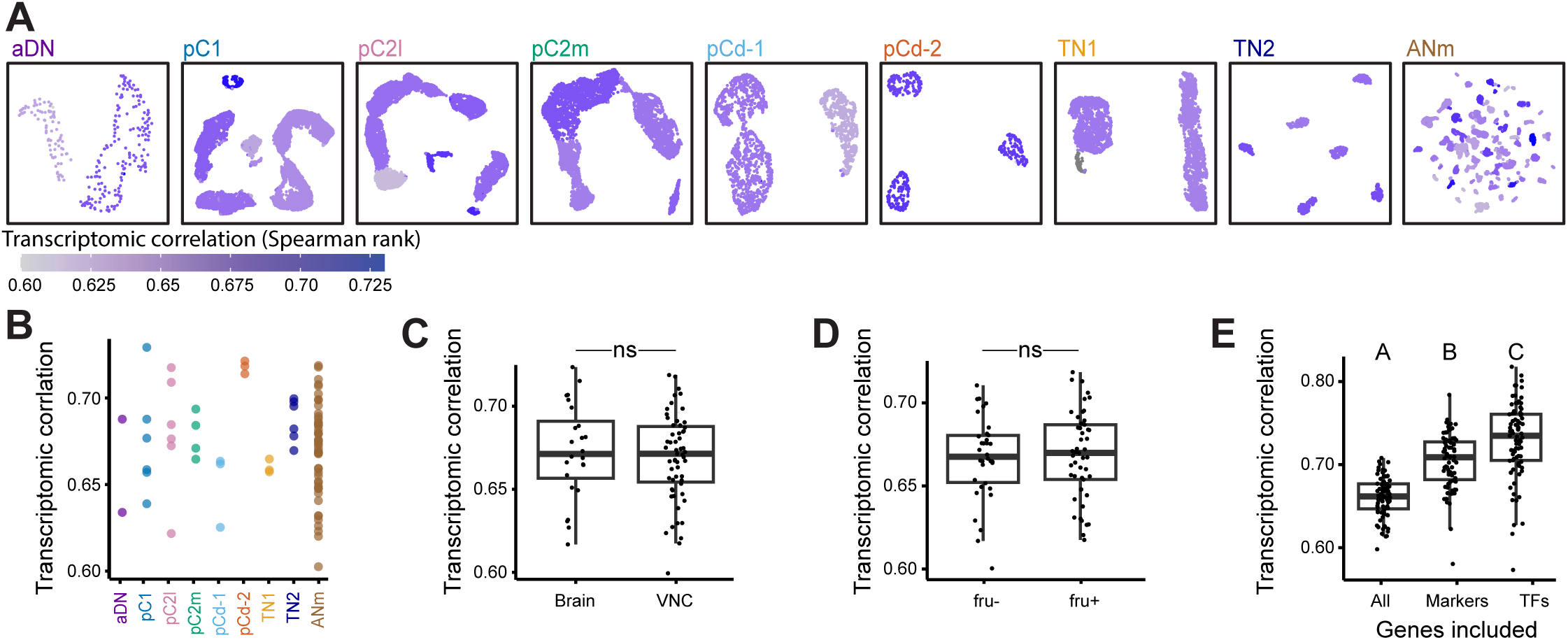
Variation across subclusters in transcriptomic conservation between *D. melanogaster* and *D. yakuba*. **(A)** UMAP representations of the subclustering of parental clusters, color-coded by transcriptomic correlation scores measured by Spearman rank correlation scores between *D. melanogaster* and *D. yakuba*. **(B)** Transcriptomic correlation scores of subclusters grouped by parental cluster; no significant difference was observed across parental clusters (Kruskal Walllis; *χ*2 = 10.793; *p* = 0.213). **(C)** Box and whisker plot showing transcriptomic correlation scores of subclusters grouped by central nervous system location, no significant difference was observed (t-test; t = 0.486, df = 39.2, *p* = 0.629). **(D)** Box and whisker plot showing transcriptomic correlation scores grouped by whether or not the subcluster expresses *fru*, no significant difference was observed (t-test; t = -0.4297, df = 72.6, *p* = 0.767). **(E)** Box and whisker plot showing transcriptomic correlation scores when including different sets of genes-all genes with median values greater than zero, only marker genes, or only transcription factors (TFs) (ANOVA; F = 82.28, df = 2, *p* < 0.001).

We examined whether the heterogeneity in transcriptomic conservation among subclusters could be caused by possible technical characteristics. First, we tested whether the across -species correlation score simply reflected how well the cell type can be molecularly defined. Cell types that are less well-defined might lead to misassignment of cell types, which may artificially cause lower correlation scores. To check this, we calculated the mean silhouette score (Tang et al., 2020), a measure of clustering quality, and found no association (**Supplemental Figure 5A**). Similarly, the correlation score was not associated with the number of cells in the subcluster ( **Supplemental Figure 5B**), or the variation across replicates **Supplemental Figure 5C**). We did find a significant positive association with the number of genes expressed within a subcluster (**Supplemental Figure 5D**). However, this did not seem to be a technical artifact because randomly downsampling the gene set to 1,000 for 1,000 iterations did not alter the association (Spearman rank correlation between the mean of bootstrap iterations and all expressed genes: r = 0.999, *p* < 0.001), raising the possibility that cell types with higher transcriptional activity may exhibit greater transcriptomic conservation. Using a generalized linear model, we found that the number of expressed genes explains 31.8% of the variance in the correlation scores among subclusters.

We further examined whether the heterogeneity could be explained by several biological characteristics. First, we asked whether specific parental clusters show greater conservation or divergence. We found that the variance of correlation scores across parental clusters was not homogenous (Bartlett test; K^2^ = 16.482; *p* = 0.0356); however, when investigating the non-homogeneity of variance by performing pairwise Bartlett tests across parental groups, there were no significant differences between any two parental clusters. We further performed a non-parametric Kruskal-Wallis test and found that the correlation scores were not different among parental clusters ( **Figure 4B**). Second, the scores also did not differ between subclusters located in the brain versus the VNC ( **Figure 4C**). Third, we tested whether they varied as a function of *fru* gene expression *fru*, given that *fru* is a key transcription factor in the development and function of sexual circuits in males (Kimura et al., 2005, 2008; Cachero et al., 2010; Rideout et al., 2010; Robinett et al., 2010) and again found no difference ( **Figure 4D**).

Together, these results suggest that transcriptomic conservation and divergence are highly cell - type-specific rather than driven by broader biological characteristics. The five subclusters with the lowest across-species correlation scores comprised four ANm subclusters (ANm_12/19/39/40) and pC2l_3. These cell types of greater transcriptomic divergence are potential candidates for functionally divergent cell types. The five most conserved subclusters include three ANm subclusters (ANm_17/23/25), pC1_6, and pC2l_6. Notably, pC1_6 also appeared more abundant in *D. yakuba*, highlighting independent evolutionary modes between transcriptomic pattern and cell type abundance.

Finally, we investigated how genes crucial to cell identity contribute to transcriptomic conservation across species. Performing Spearman rank correlations using either only marker genes or only transcription factors for Spearman rank correlations resulted in significantly higher correlation scores, with the highest observed when using only transcription factors ( **Figure 4E**). This pattern suggests stronger evolutionary constraints on the expression of genes essential for cell identity, particularly transcription factors.

### Differentially expressed genes (DEGs) across species are highly cell type-specific

We identified DEGs for each subcluster between *D. melanogaster* and *D. yakuba* and assessed their cell-type specificity. Ambient RNA in single-cell suspensions can introduce contamination, leading to false-positive signals that appear as lowly expressed genes (Macosko et al., 2015; Lun et al., 2019; Caglayan et al., 2022), artificially inflating the breadth of gene expression. Since fast-acting neurotransmitters typically show minimal co-transmission within a given cell type (Lacin et al., 2019; Shih et al., 2019; Davis et al., 2020), we used the co-expression of their marker genes (*Gad1*, *VGlut*, and *VAChT*) as a proxy for potential ambient RNA contamination (Allen et al., 2020) to explore optimal criterion of gene expression. When defining gene expression as non-zero read(s) in ≥10% of cells within a given subcluster, 20.2% (17 of 84) of subclusters appeared to release more than one neurotransmitter. In these cases, one neurotransmitter marker was expressed at a much higher level than the others, suggesting contamination. Alternatively, defining gene expression using a Bayesian mixture model by scMarc o (Chen et al., 2023), where a gene is considered expressed if it has >50% probability of being detected, yielded results that aligned more closely with expectations: 82 of 84 subclusters were shown to release at least one fast-acting neurotransmitter, with only two subclusters releasing more than one neurotransmitter, indicating limited false-negatives and false-positives. Based on this approach to filter out genes that were not expressed in either species, we further identified DEGs using a threshold of a log fold change of three and a corrected *p*-value below 0.05. Following these guidelines, 504 genes are DEGs in at least one subcluster. These genes are mostly differentially expressed in a cell type-specific manner, with many genes being differentially expressed in only a small subset of subclusters where they are expressed (**Figure 5A**), regardless of whether they are sparsely or broadly expressed ( **Figure 5B**).

**Figure 5.**
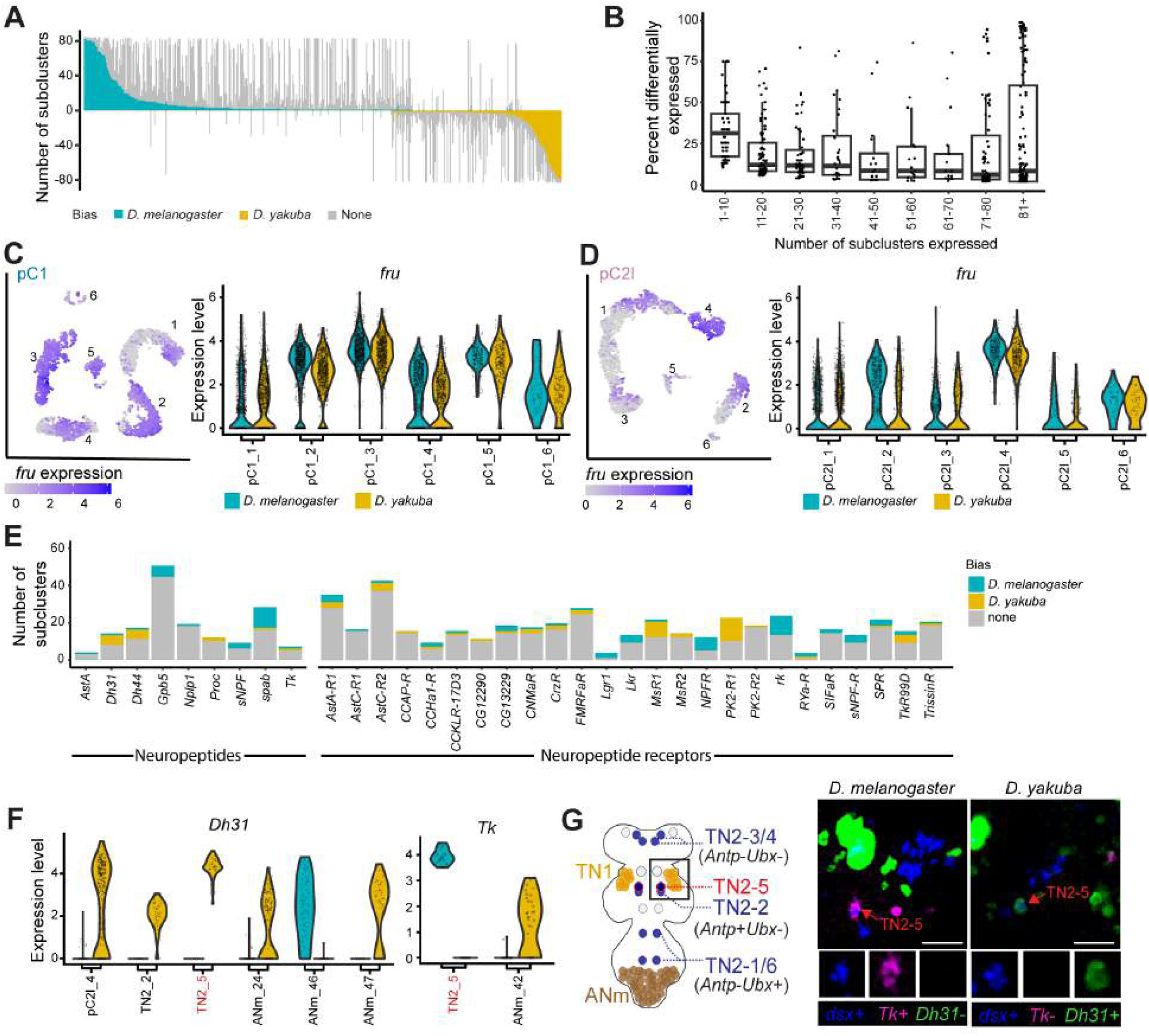
DEG analyses between *D. melanogaster* and *D. yakuba*. **(A)** Barplot showing the number of subclusters in which a gene is expressed, color-coded by the number of those subclusters in which the gene is differentially expressed and the direction of the bias. Each bar represents a single gene. **(B)** Box and whisker plot showing the percentage of subclusters in which a gene is differentially expressed, binned by the number of subclusters in which the gene is expressed. **(C)** Left: UMAP representation showing the expression of *fru* in pC1 in *D. melanogaster*. Right: Violin plots showing the expression levels of *fru* in both *D. melanogaster* and *D. yakuba* in each of the six pC1 subclusters. The expression level of *fru* is not significantly different across species in any of the subclusters. **(D)** *fru* expression in pC2l plotted similarly to panel C. **(E)** Barplot showing the number of subclusters in which neuropeptides (left) and neuropeptide receptors (right) are expressed, including only those that are differentially expressed in at least one subcluster, color-coded by the direction of bias. **(F)** Violin plots showing the expression levels of *Dh31* and *Tk* in both *D. melanogaster* and *D. yakuba* in all the subclusters where expression levels are significantly different. between the two species. **(G)** Species differences in the gene expression patterns of *Dh31* and *Tk* in the TN2-5 cell type. Left: schematic denotes the cell body location of TN2-5 and the appropriate imaging region (open black box). Right: representative confocal images of *in situ* HCR against *dsx*, *Dh31*, and *Tk* mRNA. Images presented as maximum intensity projections of z-stacks covering the cell body of TN2-5. Scale bar: 20µm.

Of the 504 DEGs, 13.3% (67 genes) exhibited higher expression in one species for certain subclusters and lower expression in others, again highlighting the cell-type specificity of gene expression evolution. The remaining genes displayed consistently higher or lower expression in one species across DEG subclusters. For these genes, we observed more DEGs with higher expression in *D. melanogaster* (60.1%, 303 genes) than in *D. yakuba* (26.6%, 134 genes). This bias cannot be explained by the percentage of reads mapped to the genome, as mapping rates were similar between species (**Supplemental Table 1**). It also cannot be readily explained by species differences in genome annotation quality, as species-specific misannotations would be expected to have a broad effect on cell types that express the gene. Here, the bias remained even when considering only genes that were not DEGs in the vast majority (>95%) of expressed subclusters (71.8% of genes consistently higher in *D. melanogaster* and 22.9% in *D. yakuba*). Therefore, this species bias might be biological.

### Across species comparison of specific genes or gene groups

#### Conserved sexual identity defined by *fru*

Changes in the spatiotemporal regulation of sex determination genes are thought to be a major mechanism driving the evolution of sex-specific traits, with many examples reported in morphological adaptations (e.g., Williams et al., 2008; Tanaka et al., 2011; Kijimoto et al., 2012; Kunte et al., 2014; Luo & Baker, 2015). However, how often such processes occur in the nervous system and contribute to behavioral evolution remains unclear. Like *dsx*, the sex determination gene *fru* plays a central role in defining the sex-specific fate of neurons and acts as a master regulator of sexual circuit development and function (Goodwin & Hobert, 2021). The *dsx+* neurons include both *fru+* and *fru-*populations with known functional differences (Rideout et al., 2007; Kimura et al., 2008; Sanders & Arbeitman, 2008; Robinette et al., 2010; Koganezawa et al., 2016; Coleman et al., 2024). For example, *fru*+ and *fru*-populations within the parental cluster pC1 exhibit behavioral specialization in courtship versus aggression (Koganezawa et al., 2016) and distinct sensory responses to sexual pheromones (Coleman et al. 2024). Consistent with *fru*’s prominent role in specifying cell fate, *fru* is a marker gene for 25 of the 84 subclusters, well above the average of 9.2 subclusters for all marker genes. Additionally, *fru* expression, both in presence/absence and quantitative levels, is highly spatially organized in UMAP space (e.g. pC1 and pC2l; **Figure 5C**, **Figure 5D**, and **Supplemental Figure 6**). While cell type-specific changes in *fru* expression might seem like a plausible evolutionary mechanism for major functional modifications, *fru* is not differentially expressed in any subcluster between *D. melanogaster* and *D. yakuba* (**Figure 5C**, **Figure and 5D**), and its expression patterns are highly similar across all four species ( **Supplemental Figure 6**). This suggests that *fru*-specified sexual identity may be highly conserved across species, and that the gain or loss of *fru* sexual identity may not be a common mode of evolutionary changes in the nervous system.

#### Conserved neuronal properties of fast-acting neurotransmitters

We annotated and compared subclusters that release the fast-acting neurotransmitters acetylcholine, GABA, and glutamate. Except for the highly diverse ANm cluster, all subclusters within each parental cluster release the same fast-acting neurotransmitter, and neurotransmitter production was consistent across all four species (**Supplemental Figure 7**). Consistent with previous studies (Zhou et al., 2014; Nojima et al., 2021; Imoto et al., 2024), pC1, pC2l, pC2m, pCd-1, TN1, and TN2 are cholinergic (*VAChT/VGAT*+), pCd-2 is GABAergic (*Gad1*+), and aDN is glutamatergic (*VGlut*+). Among the 50 ANm subclusters, 28 are cholinergic, 13 are GABAergic, seven are glutamatergic, and two lack expression of marker genes for these fast-acting neurotransmitters. Expression levels of these marker genes did not differ between *D. melanogaster* and *D. yakuba* in any subcluster.

#### Conserved neuronal properties of monoamines

Monoaminergic neurons are known to regulate sexual behaviors (Lee & Hall, 2001; Certel et al., 2007, 2010; Yilmazer et al., 2016; Zhang et al., 2016; Jois et al., 2018). To annotate monoaminergic neurons, we used *Vmat*, which encodes a monoamine vesicular transporter, as a marker gene (Greer et al., 2005). Across all four species, *Vmat* is expressed in two ANm subclusters: ANm_14 is serotonergic ( *SerT*+) and did not express maker genes of fast-acting neurotransmitters; and ANm_20 is glutamatergic ( *DAT*+) (**Supplemental Figure 8**). The serotonergic ANm_14 corresponds to a previously described neuron group that is necessary for successful copulation in *D. melanogaster* males (Yilmazer et al., 2016). Expression levels of *Vmat*, *SerT,* and *DAT* did not differ between *D. melanogaster* and *D. yakuba*.

#### Pervasive gene expression changes in neuropeptides and neuropeptide receptors

Neuropeptide signaling pathways, along with multiple terms related to synaptic connectivity and axon guidance, are among the enriched GO terms for DEGs. Here, we specifically focused on the gene expression patterns of neuropeptides and neuropeptide receptors, given their well-established functions in reproductive and courtship behaviors (e.g. Castellanos et al., 2013; Lee et al., 2015; Sellami & Veenstra, 2015; Liu et al., 2019; Wu et al., 2019) and the recognized importance of neuromodulation in behavioral evolution (Stoop, 2012; Taghert & Nitabach, 2012; Van Den Pol, 2012; Nässel & Zandawala, 2022; Jiang et al., 2024). Of the 23 neuropeptides expressed in at least one subcluster, nine are differentially expressed between *D. melanogaster* and *D. yakuba* in at least one subcluster (**Figure 5E**). Three of these genes–*Dh31*, *Dh44*, and *Tk*–have been implicated in sexual behaviors, including male courtship (Asahina et al., 2014; Shankar et al., 2015; Lee et al., 2015; Lin et al., 2022; Kim et al., 2024). Notably, all three genes are DEGs only in a subset of the subclusters where they are expressed, with expression biased toward *D. melanogaster* in some subclusters and towards *D. yakuba* in others (Figure 5D). Of the 34 neuropeptides receptors expressed in at least one subcluster, 25 are differentially expressed in at least one subcluster, with 14 whose gene expressions are biased toward different species depending on the subcluster (**Figure 5F**). These 25 genes include *CCKLR-17D3* and *SIFaR*, both of which play roles in male courtship (Wu et al., 2019, Sellami & Veenstra, 2015) and are DEGs in a small subset of expressed subclusters.

DEG analysis revealed that the cell type TN2-5 expresses the neuropeptide *Tk* in *D. melanogaster* and *Dh31* in *D. yakuba* (**Figure 5F**). We leveraged this intriguing pattern as a case for DEG validation. Based on molecular markers, we first registered TN2-5 to one of the two mesothoracic TN2 neuron pairs previously referred to as MsA (Robinette et al., 2010; **Figure 5G and Supplementary Figure 9**). Using multiplexed *in situ Hybridization Chain Reaction* (HCR; Choi et al., 2018), we then confirmed that one pair was consistently *Tk*+ and *Dh31*-in *D. melanogaster* and *Tk*- and *Dh31*+ in *D. yakuba* (the other pair was *Tk*- and *Dh31*- in both species) (**Figure 5G**). Further scRNA-seq comparisons across four species showed that *D. yakuba*, *D. santomea*, and *D. teissieri* shared the same *Tk-* and *Dh31*+ pattern in TN2-5 (**Supplementary Figure 9**). This raised the possibility that the shift in *Tk* and *Dh31* expression co-occurred at the divergence between *D. melanogaster* and the common ancestor of the other three species.

### Defining sex differences in *dsx+* cell types in *D. melanogaster*

While many *dsx*+ parental clusters are male-specific or more abundant in males, female *D. melanogaster* have about 360 *dsx*+ neurons, many of which regulate key aspects of female sexual behaviors (Lee et al., 2002; Kimura et al., 2008; Robinette et al. 2010; Rideout et al., 2010; Zhou et al., 2014; Deutsch et al., 2020; Schretter et al., 2020; Mezzera et al., 2020; Wang et al., 2020). However, the diversity of *dsx+* cell types in females and the sex differences in *dsx+* cell types–particularly in the ANm, where the majority of *dsx*+ neurons are found in females (Lee et al., 2002; Rideout et al., 2010; Deutsch et al., 2020) –remain mostly uncharacterized. To address this, we generated a scRNA-seq dataset of *dsx*+ neurons in adult female *D*. *melanogaster* to define and compare cell types between sexes. This dataset not only provides foundational knowledge of female sexual circuits and sexual dimorphism but also enables our further exploration of the relationship between species- and sex-differences (see next session).

The female data include 3,802 high quality *dsx+* cells from adult female *D. melanogaster*, resulting in 10.8x coverage. To ensure comparability, we downsampled our male *D. melanogaster* data to the same coverage and integrated them for clustering to define sex-specific and sex-biased cell types. Using the same marker genes for males (**Supplemental Figure 1**), we successfully identified each parental cluster in the integrated male/female dataset (**Figure 6A**). Consistent with previous studies (Lee et al., 2002; Rideout et al., 2010; Deutsch et al., 2020), TN1, and TN2 are male-specific, while pC1, pC2l, and pC2m are male-biased (**Figure 6A** and **6B**). We further resolved cell types by extracting each parental cluster for subclustering (**Figure 6C**) and defined sex-specific, and sex-biased cell types as those subclusters consisting of over 95% or 75% of cells from one sex, respectively. We note that this classification is based on transcriptomic patterns; it may not fully reflect the precise developmental origins. Among the 81 subclusters resolved, we classified 47 as sex-unbiased, 26 as male-specific, 2 as female-specific, 3 as male-biased, and 3 as female-biased (**Figure 6D and 6E**). The majority (89.7%) of male-specific and male-biased cell types are located in the brain, whereas all female-specific and female-biased ones are found in the ANm.

**Figure 6.**
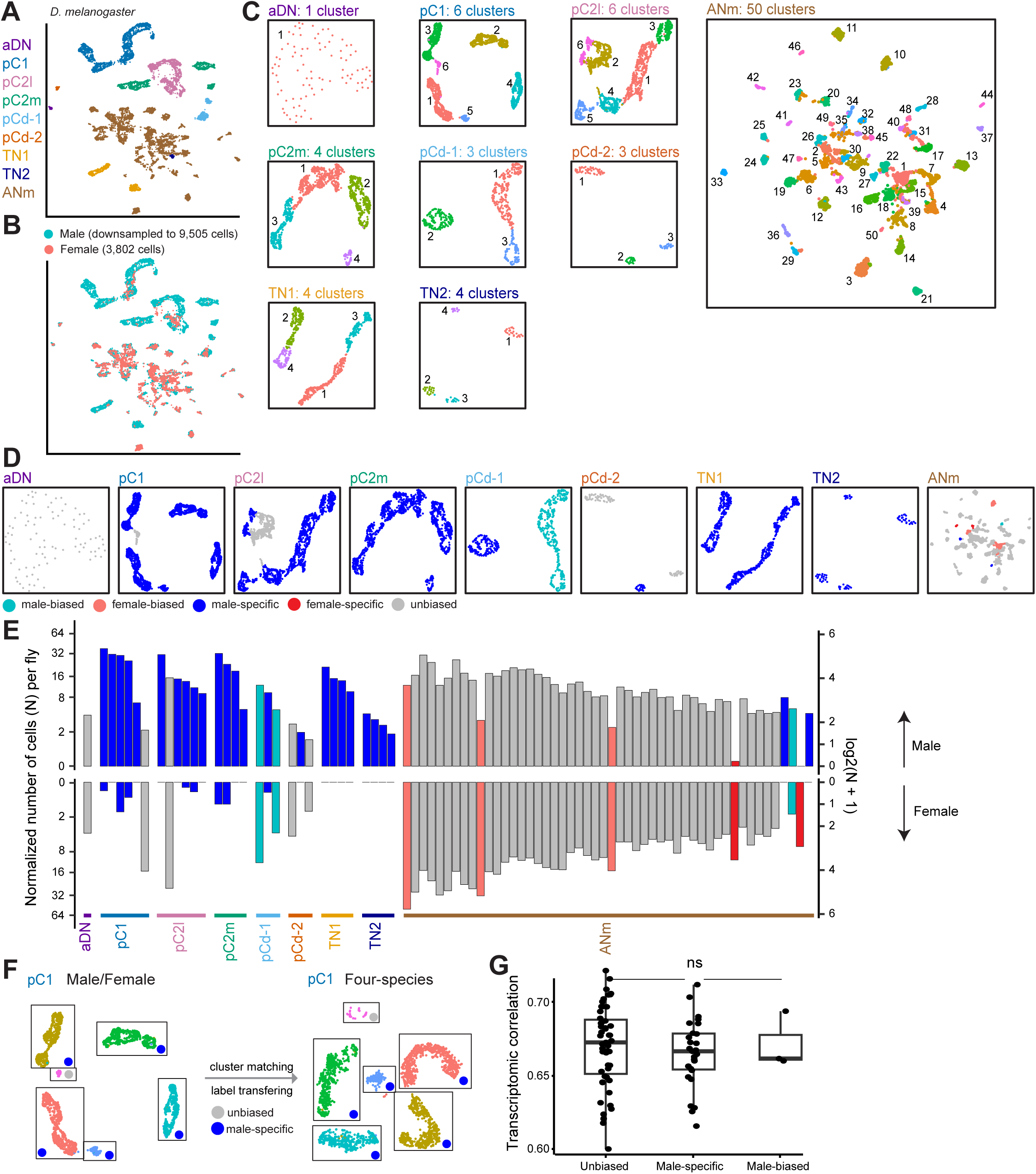
Comparison of *dsx*+ neurons across sexes in *D. melanogaster*. **(A)** UMAP representation of scRNA-seq data from *D. melanogaster* males and females, color-coded by parental cluster. **(B)** UMAP representation from panel **A** but color-coded by male and female cells. **(C)** UMAP representations of the subclustering of parental clusters shown in panel **A**. **(D)** UMAP representations of the subclusters, color-coded by sex-biases in cell number within each subcluster. **(E)** Barplot showing the normalized number of cells (N) per fly in each subcluster, separated by sex, and color-coded by sex-biases. Numbers were plotted on a log2 scale with an offset of 1 (log2(N+1)) along the Y-axis to accommodate a wide range of values and account for zeros. **(F)** Schematic representation showing our approach to match cell type identities across datasets by matching cell barcodes present in each dataset, color-coded by subcluster identity in the four species dataset. Only pC1 is shown as an example. **(G)** Box and whisker plot showing transcriptomic correlation grouped by whether the cell type is sex-unbiased across males and females, male-specific, or male-biased, no significant difference was observed (ANOVA; F = 0.0004, df = 2, *p* = 0.751).

### No clear association between sex differences and species differences

Males and females may experience different fitness consequences from the same evolutionary changes (Chapman et al., 2003; Arnqvist & Rowe, 2005; Lessells, 2006; Tregenza et al., 2006; Cox & Calsbeek, 2009). However, it remains unclear whether sex-unbiased cell types in sexual circuits are more evolutionarily constrained than sex-specific ones. Additionally, it is unknown whether sex-specific cell types are subject to stronger positive sexual selection. Therefore, we examined whether male-specific and male-biased *dsx*+ cell types are evolutionarily more divergent than the sex-unbiased ones.

To transfer the ‘male-specific’, ‘male-biased’ and ‘sex-unbiased’ labels from the male/female dataset to the four-species dataset for species comparisons, we cross-referenced the cell type identities of male cells present in both datasets (**Figure 6F** and **Supplemental Table 4**). Of the 79 subclusters in the male/female dataset (excluding the two female-specific ones), 68 had clear one-to-one match in the four-species dataset, with >80% of cells aligning to a single subcluster. The remaining 11 had more complex relationships, often due to more-to-one matches in the four-species dataset.

We tested whether male-specific cell types showed greater divergence between *D. melanogaster* and *D. yakuba* in abundance or transcriptomic patterns. Among the 26 male-specific cell types, three were either species-specific (TN1_4) or species-biased (pC1_6 and TN1_2), and one (pC2l_3) was among the five subclusters with the lowest transcriptomic correlation scores across species. However, male-specific cell types were not significantly overrepresented among species-specific or species-biased cell types (Fisher’s exact test; odds ratio = 1.338; *p* = 1) or among those with the five lowest transcriptomic correlations (odds ratio = 1.542; *p* = 1). Additionally, transcriptomic correlation scores did not differ among male-specific, male-biased, and sex-unbiased cell types (**Figure 6G**). Overall, we found no clear association between sex differences and species divergence.

### Resources for cell type-specific marker combinations and gene expression inquires

Gaining genetic access to discrete cell types is crucial for investigating the organization, function, and evolution of the nervous system. Exploiting our datasets across species and between sexes, we used *scMarco–*a graphical user interface that binarizes gene expression in scRNA-seq data using a Bayesian mixture model approach (Chen et al., 2023)*–*to identify optimal combinations of molecular markers for each cell type (**Supplemental material**). This approach informs genetic intersection strategies for cell-type-specific labeling and functional manipulations, a powerful technique that has been successfully demonstrated (Tirian & Dickson, 2017; Luan et al., 2020; Chen et al., 2023; Ye, et al., 2024).

For each subcluster defined in both the four-species dataset and the male/female dataset, we assessed individual marker genes (for genes uniquely expressed in just one cell-type) and pairs of marker genes by two key scores: (1) the clarity score (ranging from 0 to 1), which quantifies how uniquely a gene or gene pair labels a given subcluster, and (2) the bimodal score, which represents the likelihood that a gene is bimodally expressed in that subcluster. We report all genes and gene pairs with a clarity score of 1 for each of the four species individually and for male and female *D*. *melanogaster* (**Supplemental material**). We report all genes and gene pairs with clarity scores of 1, so that users can select the most convenient marker genes based on existing split-GAL4 lines or by the ease of making new lines. In the four species dataset, depending on the species, there are between 23 and 30 subclusters that can be labeled by just one gene (in addition to *dsx*), and 15 (male) or 2 (female) such subclusters in the male/female dataset. These subclusters have NA values in the columns referring to “gene 2” in the **Supplemental material**.

Users can cross-reference these marker genes or gene pairs with the *D. melanogaster* gene-specific split-GAL4 database (https://splitgal4.org/) to identify existing driver lines or design new lines for targeted genetic manipulations (Chen et al., 2023). Many split-GAL4 lines can be generated by simple genetic crosses (Li et al., 2023). Users may further intersect the split-GAL4 with *dsx^FLP^* (Rezával et al., 2014) to achieve labeling specificity as needed.

Additionally, both the four-species and the male/female datasets are publicly accessible through a Shiny app (https://apps.yenchungchen.com/dsx_neurons), an interactive web portal designed to help users identify and evaluate optimal molecular markers for cell types of interests while also enabling gene expression queries across cell types, species, and sexes. These resources will facilitate both precise genetic manipulations in *D. melanogaster* and comparative studies across species.

## Discussion

A prerequisite for understanding behavioral evolution is to decipher how changes in the molecular identity of cell types shape divergent circuit properties and behaviors. In this study, we generated a comprehensive cell type atlas of male *dsx*+ neurons across four species of *Drosophila* (**Figure 1** and **Figure 2**). Our species comparison revealed key aspects of neural circuit evolution, including cell types, transcriptomic conservation, and gene expression (**Figures 3-5**). We also generated a cell type atlas for *dsx*+ neurons in female *D. melanogaster*, defined sex differences in *dsx+* cell types, and tested whether male-specific cell types exhibit distinct evolutionary patterns ( **Figure 6**). Finally, we systematically identified combinations of molecular markers that uniquely label each cell type, enabling precise cell type-specific functional characterization in future studies.

By isolating relevant neurons associated with specific behaviors of rapid evolution, our study provided the previously lacking granularity in cell type evolution. Among the 84 defined cell types, ranging from one to about 23 neurons per hemisphere, we identified all but two across all four *Drosophila* species. We note that the appropriate level of granularity in defining cell types is an open question (Zeng, 2022; Özel & Desplan, 2025). Here, we refined cell types by first identifying biologically meaningful parental clusters and then iteratively subdividing each until further division no longer yielded strong marker genes. More cell types may still be embedded in our dataset. For instance, several pC1 and pC2l subclusters contain both *fru*+ and *fru*-populations, suggesting potential heterogeneity. Clustering strategies, such as species selected for data integration, may also influence the precise outcome of cell type identification. Nevertheless, our study provides a necessary baseline for investigating the cellular basis of behavioral evolution. The high-resolution datasets generated here allow continued exploration tailored to specific research questions and cell types of interest, such as adjusting clustering parameters, performing clustering on unintegrated or differently integrated datasets, combining scRNA-seq data of earlier developmental stages, and incorporating additional neuronal modalities such as neuron anatomy informed by light microscopy and EM connectome.

In contrast to the overall conservation in the presence and absence of cell types, species differences in cell type abundance appear to be common, accompanied with widespread transcriptomic changes in a highly cell-type-specific manner. 12 of 84 cell types, spanning diverse functional populations, potentially vary in abundance between *D. melanogaster* and *D. yakuba*. At the transcriptional level, we observed substantial variation in transcriptomic conservation even among cell types within the same parental cluster. This heterogeneity also aligns with the cell-type-specificity of DEGs, as most of the 490 DEGs we identified were differentially expressed in only a small subset of cell types across species. Interestingly, the heterogeneity among cell types cannot be readily explained by various features we examined, such as cluster size, parental cluster identity, or sex specificity. These findings together suggest that the evolutionary processes shaping transcriptomic divergence, including “developmental system drift” and adaptive changes of new functions, operate in a highly cell-type-specific manner.

At the level of individual cell types, our study revealed conserved and evolvable aspects of gene expression in sexual circuits. Surprisingly, despite long-standing appreciation of *cis*-regulatory changes of master regulators as key drivers of phenotypic evolution (Wittkopp & Kalay, 2012), *fru*–a central orchestrator of male sexual circuits (Goodwin & Hobert, 2021) as well as a previously implicated genetic locus of species divergence (Gleason & Ritchie, 2004; Lagisz et al., 2012; Parker et al., 2014) –did not stand out as an obvious player. Both the composition of *fru*+ and *fru*-populations in *dsx*+ neurons and the quantitative expression of *fru* across cell types appear highly conserved. (we note that a substantial proportion of *fru*+ neurons do not express *dsx*, thus species differences in *fru* expression that were not investigated here may exist). This observation aligns with recent work revealing a conserved functional subdivision of P1 neurons–a subset of pC1 neurons that acts as a core node of the male courtship circuit–based on *fru* expression (Coleman et al., 2024). Similarly, the conservation of *dsx*+ cell types across species, along with the absence of quantitative species differences in *dsx* expression levels, suggests that *dsx*-defined sexual identity is largely conserved. Other conserved features include greater transcriptomic conservation for genes crucial to cell identity (i.e., transcriptional factors and molecular marker genes) and a fully conserved pattern of neurotransmitter identities (i.e., fast-acting neurotransmitters and monoamines) across all *dsx+* cell types in all four species. Given these conserved patterns, the widespread gene expression differences observed between *D. melanogaster* and *D. yakuba–*representing 10.2% of all genes expressed in the dataset–likely arise disproportionately from genes outside these fundamental properties. Particularly, genes involved in neuropeptide signaling, which have been broadly associated with behavioral evolution ( Katz, 2011; Schoofs et al., 2017), are significantly overrepresented in DEGs. Strikingly, 39.1% of neuropeptide genes and 73.5% neuropeptide receptor genes expressed in *dsx*+ neurons, including five with established roles in regulating sexual behaviors, are differentially expressed in at least one cell type. These findings highlight the remarkably flexible and modular nature of their gene expression patterns in evolution. Together, our study revealed a dynamic evolutionary landscape of widespread cell-type-specific modifications built upon a largely conserved organization of male sexual circuits in cell types, sexual identity, and neurotransmitter identity.

The profound functional diversity among cell types in the nervous system, coupled with the highly cell-type-specific mode of evolution, urgently calls for cell type-specific genetic tools for functional characterization. This need becomes even more critical for studying evolutionary changes, as comparing truly homologous neurons or neuronal groups across species is indispensable for appropriate functional interpretation. While transferring GAL4 and split-GAL4 enhancer drivers that label specific neurons in *D. melanogaster* into closely related species has proven to be a fruitful approach in comparative studies (e.g., Stern et al., 2017; Seeholzer et al., 2018; Ding et al., 2019; Luan et al., 2020; Ohashi et al., 2023; Ye, et al., 2024; Li et al., 2024; Coleman et al., 2024), this approach can be constrained by factors such as species differences in enhancer activity and the lack of well-characterized reagents in *D. melanogaster*. Unlike enhancer-based genetic tools, gene-specific tools that leverage native regulatory elements have been shown to reliably recapitulate the expression patterns of targeted genes (Diao et al., 2015; Lee et al., 2018; Chen et al., 2023). By identifying molecular markers and marker combinations that represent specific cell types, our scRNA-seq datasets provide a roadmap for designing gene-specific genetic tools that target individual cell types within and across species.

Finally, we have uncovered many potential evolutionary changes, including alterations in cell type abundance, cell types exhibiting pronounced transcriptomic divergence between species, and genes with species-specific expression patterns, inviting future validation and functional characterization. As a case of validation, we demonstrated that the single-neuron cell type TN2-5 has undergone an evolutionary switch in neuropeptide identity—expressing *Tk* in *D. melanogaster* and expressing *Dh31* in *D. yakuba*. While the function of TN2-5 is currently uncharacterized, its male-specificity and mesothoracic location suggest that it might be involved in generating male-specific wing behavior, such as courtship song and agnostic song in aggression (Jonsson et al., 2011). Interestingly, in *D. melanogaster*, *Tk* signaling suppresses courtship and enhances aggression (Asahina et al., 2014; Shankar et al., 2015); while gut - derived *Dh31* signaling promotes courtship (Lin et al., 2022). Future research may uncover the behavioral role of TN2-5 and the phenotypic consequences of this evolutionary switch in neuropeptide identity, now made possible by identifying molecular markers for cell-type-specific manipulation. Given the widespread evolutionary turnover in the expression of neuropeptide signaling genes, a broadly relevant question is whether the behavioral states associated with these genes have diverged across species or if the gene-behavioral state associations remain conserved while evolutionary changes in gene expression have repurposed the same cell type to associate with a distinct behavioral state.

In conclusion, using *dsx* as a molecular handle, we characterized the cellular and molecular diversity of sexual circuits with unprecedented detail both within and across species. Our findings on the conservation and divergence of cell types and gene expression provide a crucial foundation for understanding the fundamental principles governing neural circuit organization and evolution in behavioral adaptations.

## Methods

### Fly strains and husbandry

We housed all flies on a cornmeal-agar-yeast medium (Fly Food B, Bloomington Recipe, Lab Express) in a humidified incubator at 23°C on a 12 hour light and dark cycle. We list the genotypes of fly lines used in this study in **Supplemental Table 5**.

### Generation of genetic reagents

We digested the plasmid pBac {3XP3::EYFP, attP} (Stern et al., 2017) with NaeI and inserted a DNA cassette containing 10XUAS-nls::tdTomato-SV40, PCR amplified from the plasmid pJFRC105-UAS-nls::tdTomato (gift of Gerry Rubin’s lab, Janelia) into the NaeI-digested pBac {3XP3::EYFP, attP} by Gibson Assembly. We injected the resulting plasmid into *D. teissieri* wild-type flies (DSSC # 14021-0257.01) to generate the transgenic line pBac {UAS-nls::tdTomato, 3XP3::YFP}. We also injected the plasmid pJFRC105-10XUAS-nls::tdTomato into the *D. santomea* line 2253 (Stern et al., 2017) to create the transgenic line w; ; pJFRC105-10XUAS-nls::tdTomato (2283).

### Dissection, cell dissociation, library preparation, and sequencing

We genetically labeled *dsx*+ neurons by crossing females carrying *dsx*-GAL4 (GAL4 was introduced into the endogenous *dsx* locus by CRISPR/Cas9-mediated genome editing) (Ye et al., 2024) with males carrying UAS-nls::tdTomato. The resultant *dsx-*GAL4>UAS-nls::tdTomato males were sorted within 12 hours of eclosion and housed in groups of 15–20 until dissection at 4–8 days old. We dissected the CNS (brain and ventral nerve cord) from approximately 100 flies per sample. Our cell dissociation method was adapted from previously described protocols (Li et al., 2022). In short, we dissected the flies in ice-cold Schneider’s medium for one hour before transferring all tissue into 500 ul of dispase (1mg/1ml Liberase DH dissolved in Schneider’s medium). We digested the tissue in 500 µL dispase (1 mg/mL Liberase DH in Schneider’s medium) for 45 minutes at room temperature, then washed it three times with ice-cold Schneider’s medium. We dissociated the tissue in 500 µL ice-cold 1X PBS with 2% BSA by pipetting it 100 times with a pipette and 20 times with a 25-gauge needle, both coated with FBS to prevent cell adhesion. We transferred the solution into a flow cytometry tube before sorting out nls::tdTomato+ ( *dsx*+ neurons) cells with a FACS Aria Cell Sorter (using a 100 µm nozzle and 1.25 drop precision). To verify the yield, we counted the number of nls::tdTomato+ cells using a disposable hemocytometer and a Leica M165 fluorescent scope. Upon confirmation, we transported the cells to the Center for Applied Genomics at the Children’s Hospital of Philadelphia’s Research Institute where all library preparation and sequencing was performed using the 10x Genomics Single Cell 3’ Library and Gel Bead Kit following the manufacturer’s recommendation protocol and the libraries were sequenced on an Illumina NovaSeq 6000. The amount of time between the start of cell dissociation and the start of cell sorting was about 70 min, sorting the cells lasted about 30 min, and the time between the end of cell sorting and the start of the 10x Genomics protocol was about 30 min. We prepared two replicates for both *D. melanogaster* and *D. yakuba* (in addition to using one published replicate per species from Ye et al., 2024) and one sample each for *D. teissieri* and *D. santomea*.

### scRNA-seq data mapping and filtering

We used the CellRanger software (v7.1.0) “count” command to map sequencing reads to *Drosophila melanogaster* (Assembly GCA_000001215.4), *Drosophila yakuba* (Assembly GCA_016746365.2), *Drosophila santomea* (GCA_016746245.2), and *Drosophila teissieri* (GCA_016746235.2) genomes accordingly. We analyzed expression matrices with Seurat v5.0.1 (Hao et al., 2024) and applied default parameters for all Seurat functions unless otherwise specified. To ensure high-quality single-cell data, we filtered the datasets by removing cells with greater than 15% mitochondrial transcripts, fewer than 200 genes detected, or expressing more than 4000 unique genes. To identify orthologous genes between species, we performed three reciprocal best hits BLAST analyses, pairing *D. yakuba*, *D. santomea*, and *D. teissieri* with *D. melanogaster* using default settings (Camacho et al., 2009). We identified 11,739 orthologous genes shared among all four species and retained only these genes in the datasets, including the dataset of *D. melanogaster* males and females (described below).

### scRNA-seq data clustering and integration

We used the functions “NormalizeData” and “ScaleData” to normalize and scale the individual species’ datasets, followed by principal components analyses (PCA). We determined the number of statistically significant principal components using Jackstraw analysis. We used the functions “runUMAP”, “FindNeighbors”, and “FindClusters” to reduce the dimensionality and perform clustering. We assessed clustering quality using the R package scclustereval v0.0.0.9 (Tang et al., 2020). After clustering, we removed clusters where the majority of cells lacked expression of *dsx* or *elav* (a neuronal marker gene), retaining a total of 17,168 *D. melanogaster*, 18,067 *D. yakuba*, 6,344 *D. teissieri*, and 5,343 *D. santomea* cells. See **Supplemental Table 1** for a summary of all sequencing and filtering data.

We initially analyzed each species data separately to confirm the presence of all nine parental clusters in each species (see below). After identifying the nine parental clusters in each species, we integrated the datasets across species using the Seurat functions “SelectIntegrationFeatures”, “FindIntegrationAnchors”, and “IntegrateData”. Following integration, we performed PCA and Jackstraw analysis, reduced dimensionality, and conducted clustering. We confirmed the presence of all nine parental clusters in the integrated dataset and extracted each parental cluster into a separate Seurat fil e for further clustering analyses using the same protocol described above.

### Parental clusters annotations

To identify each parental cluster, we first classified neurons as either from the brain or the VNC based on the expression of *tsh* (a VNC marker gene; Röder et al., 1992) and the four Hox genes *Antennapedia* (*Antp*), *Ultrabithorax* (*Ubx*), *abdominal-A* (*abd-A*), and *Abdominal-B* (*Abd-B*), which follow an anterior-to-posterior expression pattern in the VNC (Allen et al., 2020). In the brain, we considered the expected size for each parental cluster and identified good marker genes for clusters defined in UMAP space. We then used split-GAL4, pairing a *dsx* hemi-driver with a hemi-driver for each marker gene, to drive the expression of fluorescent proteins (GFP or tdTomato) in *D. melanogaster*. We performed immunostaining against the fluorescent proteins to determine the *dsx*+ neurons labeled by each marker gene. These maker genes included *tey*, *Optix*, *TfAP-2*, *sv, unc-4, Gad1*, whose intersection with *dsx* labeled aDN, pC1, pC2l, pC2m, pCd-1, and pCd-2, respectively. In the VNC, the marker gene *TfAP-2* labeled TN1 neurons (Ye, et al., 2024). We further identified neurons that were *Antp*+ and/or *Ubx*+ but *abd-A*- and *Abd-B*-, hypothesizing that these were TN2 given that neurons in the ANm should express one or both of *abd-A* and *Abd-B*. A split-GAL4 intersection between *ara*–a marker gene for these neurons–and *dsx* confirmed their identity as TN2. We scored all VNC clusters that were not TN1 or TN2 but expressed *abd-A* and/or *Abd-B* as ANm. Immunostaining was performed as previously described (Ye et al., 2024) and imaged on a Leica 350 DMi8 microscope with a TCS SP8 Confocal System at 40x with optical sections at 0.4-0.8 µm intervals. All the genes mentioned above were shared marker genes identified across independent clustering analyses for each of the four species, permitting us to transfer parental cluster identities from *D. melanogaster* to *D. yakuba*, *D. santomea*, and *D. teissieri*. Beyond relying on these shared marker genes, we further validated the transferability of our annotations by measuring the similarity of parental clusters across species using MetaNeighbor (Crow et al., 2018) to calculate the mean AUROC scores for each pairwise comparison.

### Identification of marker genes

To identify marker genes for all parental clusters and subclusters, we used the Seurat function “FindAllMarkers” with the “assay” parameter set to “RNA”. This function employs Wilcoxon signed-rank tests to identify marker genes. We performed this analysis for all four species combined as well as for each species individually.

### Designating parental cluster- or subcluster-level gene expression

We defined a gene as expressed in a given subcluster if the *scMarco* (Chen et al., 2023) bimodal score for that gene in the subcluster exceeded 0.5. Using this criterion, we annotated hemilineage markers, *fru*, neurotransmitter markers, neuropeptides, and neuropeptide receptors. We also quantified the number of genes expressed in each subcluster and identified genes to include in our DEG analyses (see below).

### Predicting hemilineage identity

We used previously identified hemilineage markers (Bossing et al., 1996; Schmid et al., 1999; Lacin et al., 2019; Allen et al., 2020) to annotate the hemilineage origins of ANm subclusters. We considered a marker gene expressed in a given subcluster if the scMarco bimodal score exceeded 0.5 for *D. melanogaster* cells within the integrated dataset. See **Supplemental Table 2** for a full list of hemilineage annotations and the marker genes used.

### Defining species differences in cell number within subclusters between *D. melanogaster* and *D. yakuba*

For each species, we manually quantified the number of cells in each *dsx+* parental cluster in *dsx*-GAL4>UAS-myrGFP males and females by immunostaining as previously described (Ye et al., 2024). We counted cells from confocal images using the Fiji/ImageJ plugin “Cell Counter.” We excluded the ANm cluster due to the high density of *dsx+* neurons, which made precise counting challenging, and therefore assumed equal numbers of ANm *dsx*+ cells in species comparison. Using these data, we scaled the cell count for each cell type to align the size of each parental cluster with immunostaining measurements. After scaling, we applied a Bayesian modeling approach with the Python package *scCODA* v0.1.2 (Büttner et al., 2021) and default parameters to identify species-biased subclusters.

### Spearman rank correlation analysis across species

To assess transcriptomic conservation between *D. melanogaster* and *D. yakuba* subclusters, we calculated Spearman rank correlations based on the median expression levels of genes with non-zero median expression. We performed this analysis using three gene sets: all genes, all marker genes, and transcription factors. For each subcluster, we also conducted 1,000 iterations of randomly downsampling the gene set to 1,000 genes (selected from genes with non-zero median expression in at least one species) and calculated the Spearman rank correlation between *D. melanogaster* and *D. yakuba*. Additionally, we computed the Spearman rank correlation for each subcluster between replicates within each species, considering all genes with non-zero median expression values. When grouping Spearman rank correlation scores by parental cluster, we found that although the data were normally distributed (Shapiro-Wilk, W = 0.985, *p* = 0.462), the variance across parental clusters was not homogenous (Bartlett test; K^2^ = 16.482; *p* = 0.0356). Therefore, we conducted a nonparametric Kruskal Wallis test. Furthermore, we conducted pairwise Bartlett tests across parental clusters but found no significant differences after correcting for multiple comparisons via a Bonferroni adjustment.

### DEG analysis between *D. melanogaster* and *D. yakuba*

To identify differentially expressed genes (DEGs) between *D. melanogaster* and *D. yakuba*, we used the Seurat function “FindAllMarkers” with the “assay” parameter set to “RNA” and the active Ident set to species after subsetting out only the *D. melanogaster* and *D. yakuba* cells from the full integrated dataset. For each subcluster, we included genes expressed in at least one of the two species defined by scMarco to perform DEG analysis. For each subcluster, we compared the expression of each gene in *D. melanogaster* cells to its expression in *D. yakuba* cells. To ensure that we took a relatively conservative approach, we classified genes as differentially expressed only if they met a log fold change threshold greater than three and had Bonferroni-corrected *p*-values less than 0.05. We performed Gene Ontology (GO) term enrichment analysis using PANTHER (Mi et al., 2015), looking for enriched “biological process” terms among the DEGs against all genes e expressed in at least one subcluster.

### Validating differential gene expression of *Tk* and *Dh31* in TN2-5 neurons between species

To determine the identity of TN2-5 neurons, we performed immunostaining against *Antp* (DHSB 8C11, 1:20) and *Ubx* (DHSB FP3.38, 1:20) in *dsx*-GAL4>UAS-myrGFP male flies (anti-GFP, Abcam #13970, 1:600). We identified the two prothoracic TN2 neurons as *Antp-* and *Ubx-*, corresponding to TN2-3 and TN2-4; the two metathoracic ones as *Antp*- and *Ubx*+, corresponding to TN2-1 and TN2-6; and the two mesothoracic ones as *Antp*+ and *Ubx*-, corresponding to TN2-2 and TN2-5. To validate the DEG genes *Tk* and *Dh31*, we performed multiplexed HCR with three probe sets detecting *dsx*, *Tk*, and *Dh31* mRNA in male flies. For *Tk* and *Dh31*, *D. melanogaster* sequences were provided for probe design (Molecular Instrument), with the request of designing probes against conserved gene regions. The same probe set was used for *D. melanogaster* and *D. yakuba*, producing robust and unambiguous signals, albeit a bit weaker in *D. yakuba*. HCR was performed following a previously described protocol (Duckhorn et al., 2022). See **Supplementary Table 5** for probe and hairpin information.

### Generating scRNA-seq data of *dsx*+ neurons in female *D. melanogaster*

To prepare the female dataset, we followed the same dissection, cell dissociation, library preparation, sequencing, data mapping, and filtering protocols as described above, resulting in 3,802 high quality *dsx*+ cells. Because males have approximately 2.5 times the number of *dsx*+ neurons as females, we randomly downsampled the male *D. melanogaster* dataset to 2.5 times the number of female cells (9,502 cells). We then integrated the downsampled male dataset with the female dataset and performed clustering using the protocol described above. Using the same marker genes as before, we identified all nine parental clusters in the male/female integrated dataset. We extracted each parental cluster individually for subclustering.

To match subclusters between our two datasets (four-species versus male/female *D. melanogaster*), we cross-referenced the subcluster identities of male cells present in both datasets using their unique barcode IDs. We considered subclusters to have a one-to-one match if over 80% of the cells from a given subcluster in the male/female dataset aligned to a single subcluster from the four-species dataset. When subclusters did not have a clear one-to-one match, we considered subclusters from the male/female dataset to contain all subclusters from the four-species dataset with over 10% matching cells. See **Supplemental Table 4**.

### Defining sex differences in cell number

Since lacked biological replicates for female *D. melanogaster*, we could not use *scCODA* to identify sex-specific or sex-biased subclusters. Instead, we defined sex-specific subclusters as those consisting of over 95% cells from one sex and sex-biased subclusters as those with over 75% cells from one sex.

### Using marker gene lists to identify genes that label subclusters

For both the four-species and the male/female datasets, we report pairs of genes that uniquely label each subcluster. Although we report all pairs of genes with clarity scores of 1, we encourage users to be selective when choosing genes by using genes with the highest possible percentage of cells expressing that gene and checking gene expression patterns in scMarco.

## Supporting information

SData1

SData2

SData3

## Acknowledgement

We thank Lihua Li and Hongjie Li for their valuable input during the development of our scRNA-seq pipeline. We thank Troy Shirangi’s lab for contributing to sample dissection. We thank Nancy Zhang for valuable feedback on statistical analysis. We are immensely grateful for the generosity of the fly research community. Particularly, we thank Haluk Lacin, Yufeng Pan, Chris Doe, Gary Struhl, Joshua Kavaler, Nobert Perrimon, Ben Ewen-Campen, Ruth Lehmann, and Jing Wang for generously sharing genetic reagents and antibodies of marker genes. Although many experiments using these resources were not directly presented here, they have greatly informed our initial identification of *dsx*+ parental clusters before we took a split-GAL4 strategy for final validation. We also thank Rory Coleman, Troy Shirangi, Dawn Chen, Stephen Goodwin, Megan Goodwin, and Aaron Allen for helpful comments on the manuscript. We thank Jennifer Jakubowski, Shifu Tian, and Andrew Morschauser for their help with FACS. We thank Molly Gallagher and Diana Slater at the Center for Applied Genomics at the Children’s Hospital of Philadelphia’s Research Institute for help with library preparation and sequencing. Y.D. was supported by NIH grant R35GM142678. Y.-C.C. was supported by New York University (MacCracken Fellowship), by a NYSTEM institutional training grant (Contract #C322560GG), and by a Scholarship to Study Abroad from the Ministry of Education, Taiwan. Y.-C.D.C. is supported by the NIH National Eye Institute (K99EY035757).

## Author contribution

J.T.W and Y.D. conceived the study and wrote the paper. I.P.J. and H.G. performed *in situ* HCR and immunostaining and collected the confocal imaging data. J.T.W. generated the scRNA-seq data and performed analyses. Y.D. generated genetic reagents. Y.-C.D.C. and Y.-C.C. generated the resources for cell type-specific marker combinations.

