## Supplemental figures and tables for "High-resolution comparative single-cell transcriptomics of *doublesex*-expressing neurons reveals evolutionary conservation and diversity of sexual circuits in *Drosophila*"

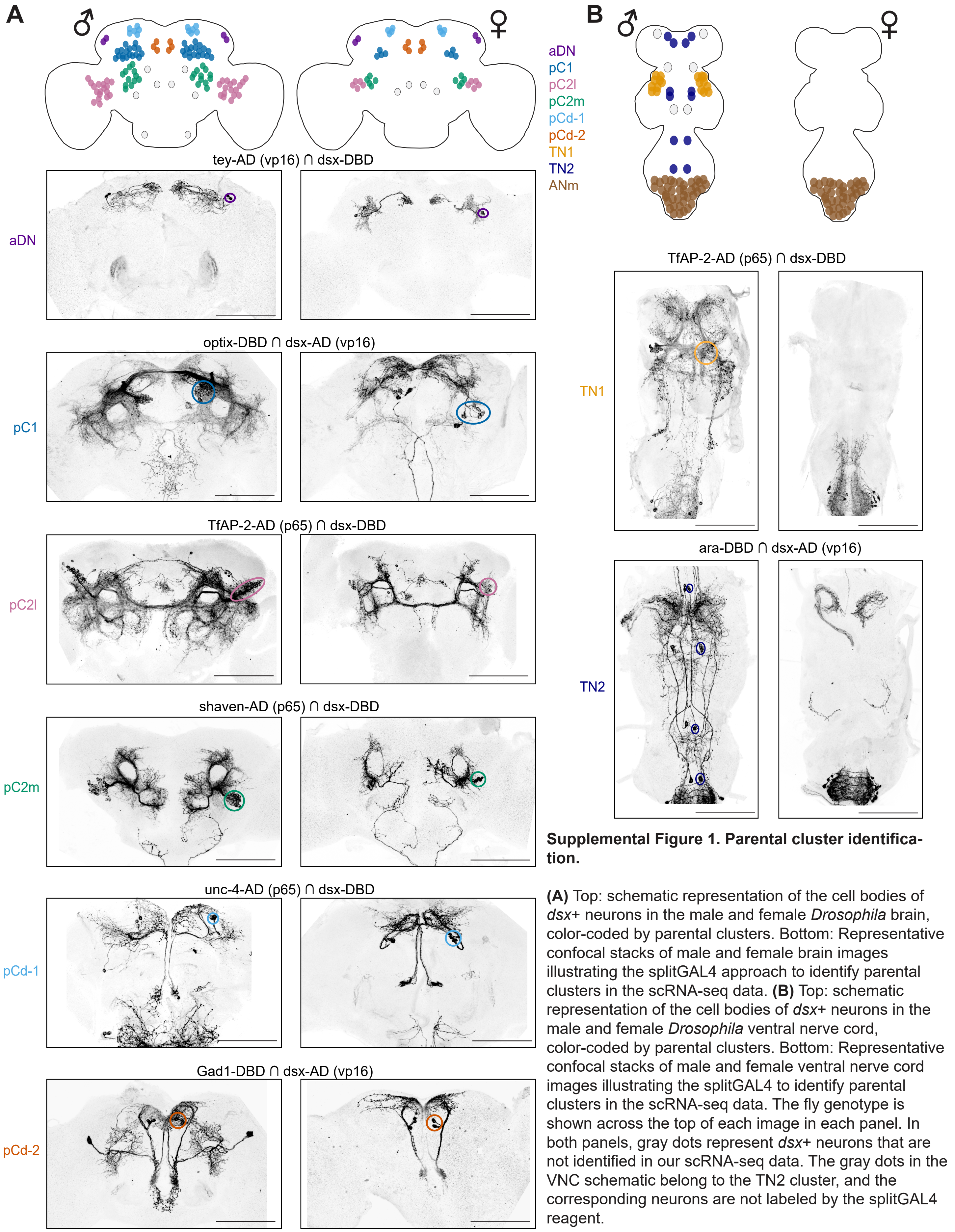

**Supplemental Figure 1. Parental cluster identification.**

**(A)** Top: schematic representation of the cell bodies of *dsx*+ neurons in the male and female *Drosophila* brain, color-coded by parental clusters. Bottom: Representative confocal stacks of male and female brain images illustrating the splitGAL4 approach to identify parental clusters in the scRNA-seq data. **(B)** Top: schematic representation of the cell bodies of *dsx*+ neurons in the male and female *Drosophila* ventral nerve cord, color-coded by parental clusters. Bottom: Representative confocal stacks of male and female ventral nerve cord images illustrating the splitGAL4 to identify parental clusters in the scRNA-seq data. The fly genotype is shown across the top of each image in each panel. In both panels, gray dots represent *dsx*+ neurons that are not identified in our scRNA-seq data. The gray dots in the VNC schematic belong to the TN2 cluster, and the corresponding neurons are not labeled by the splitGAL4 reagent.

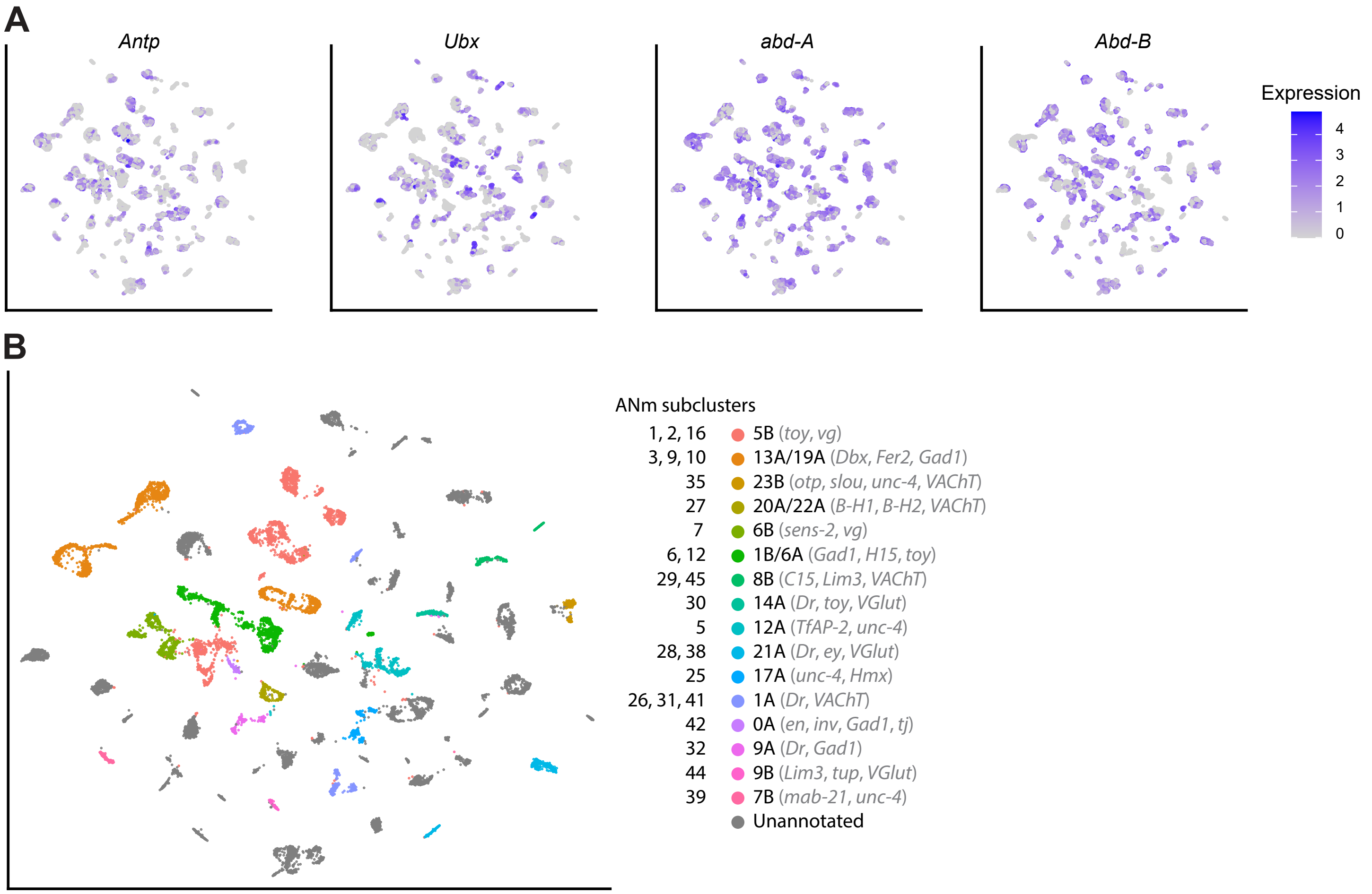

**Supplemental Figure 2. Gene expression patterns in the ANm of the integrated four-species dataset.**

**(A)** Gene expression of four Hox genes in the ANm. **(B)** UMAP representation showing ANm subclusters, color-coded by their hemilineage origin as identified using the marker genes shown in grey.

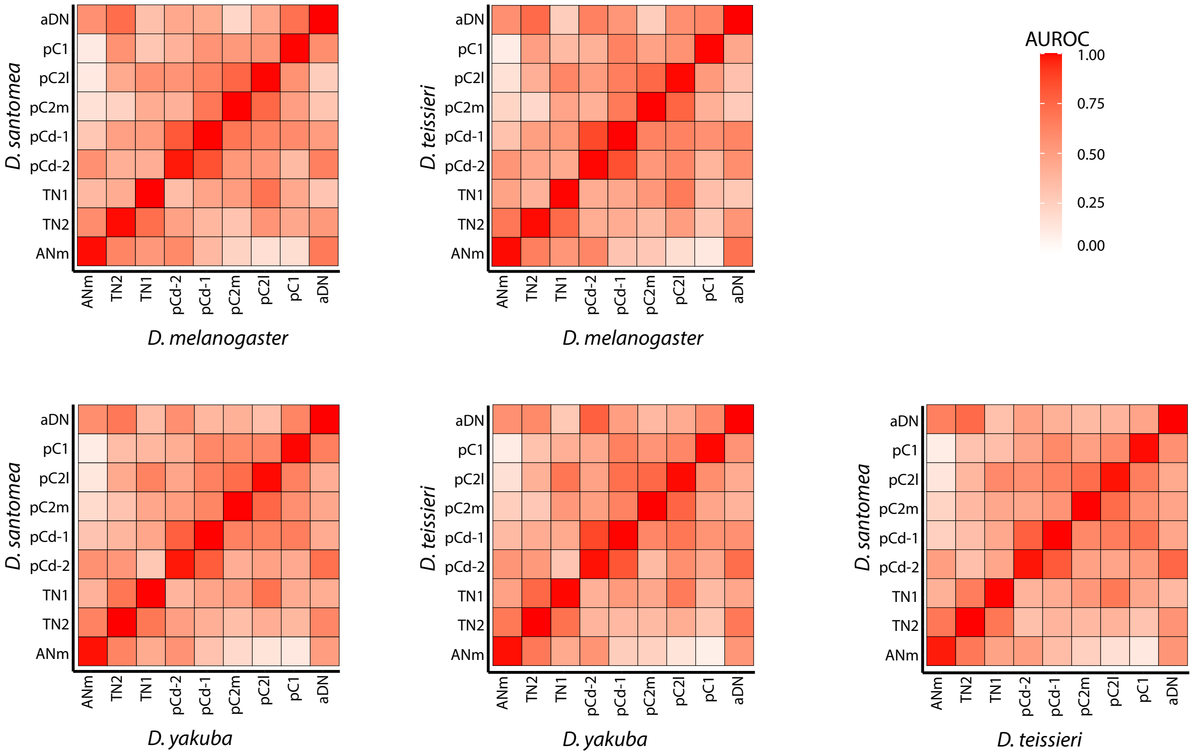

**Supplemental Figure 3**

Heatmaps showing the similarity (AUROC) of parental clusters between species.

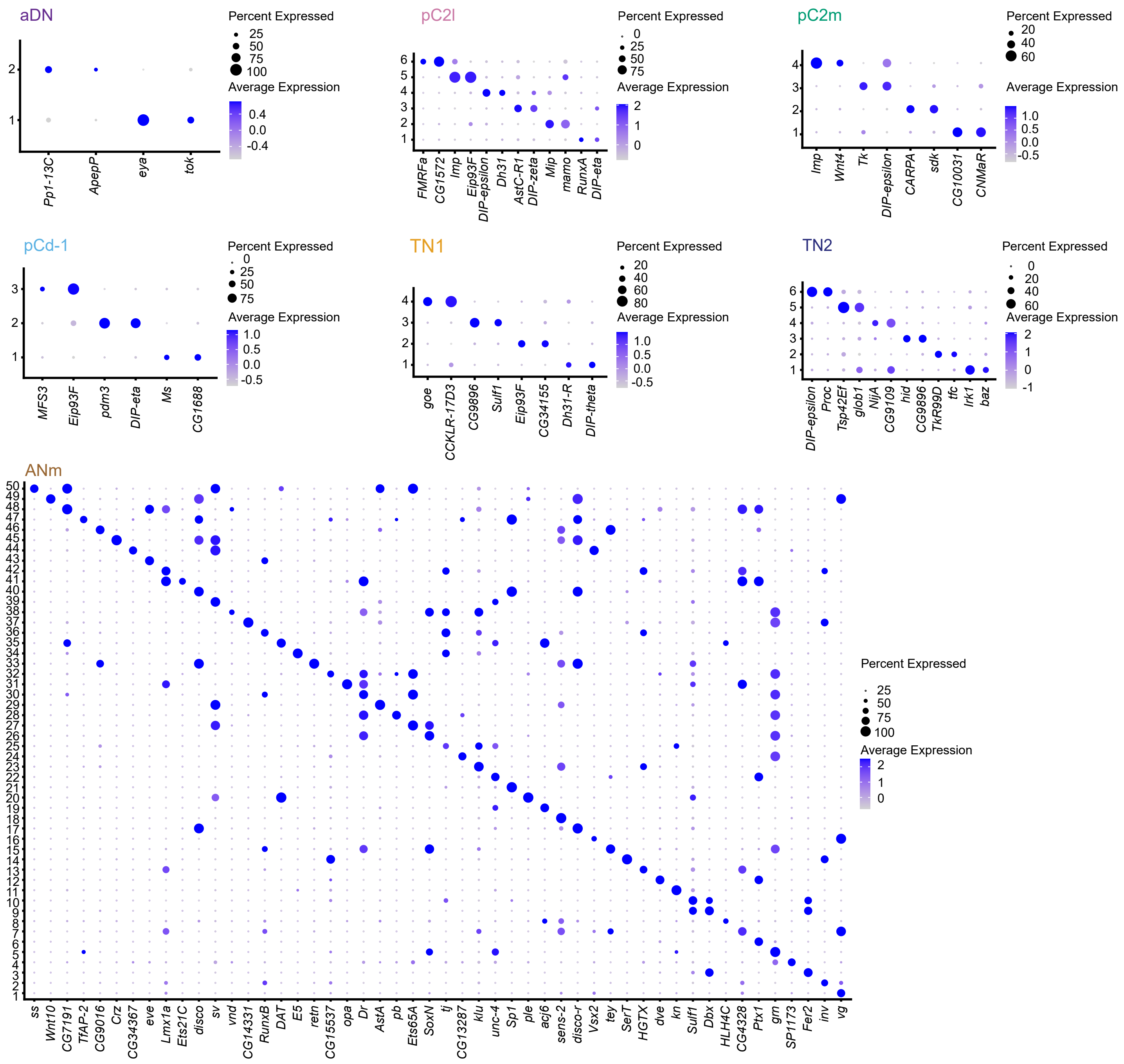

**Supplemental Figure 4**

Dot plots showing the average expression and percentage of cells expressing marker genes for subclusters grouped by parental cluster. Data are from the integrated four-species dataset.

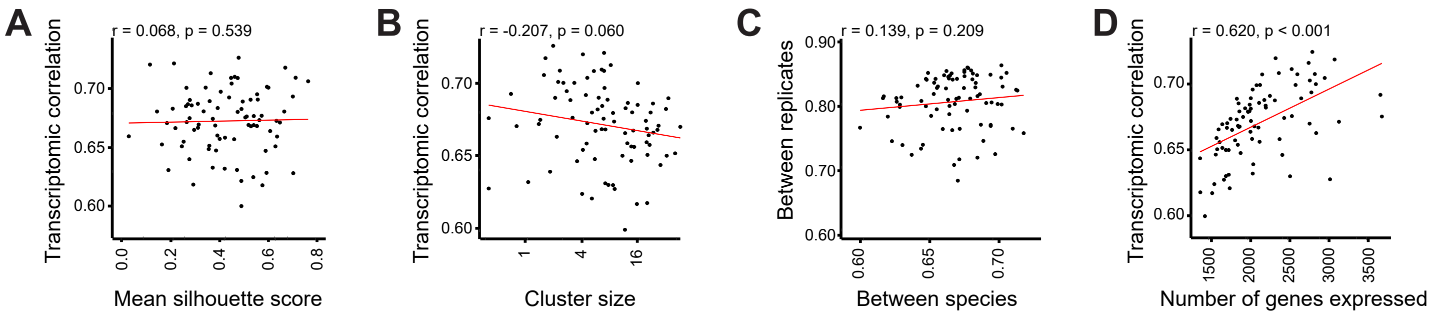

**Supplemental Figure 5. Transcriptomic conservation and technical characteristics.**

**(A)** Scatterplot showing transcriptomic correlation scores over the average silhouette score (a measure of clustering quality) for each subcluster. **(B)** Scatterplot showing transcriptomic correlation scores over the scaled subcluster size (scaled to reflect neuron counts) for each subcluster. **(C)** Scatterplot showing the average transcriptomic correlation between replicates within each species tover the ranscriptomic correlation scores across species. **(D)** Scatterplot showing transcriptomic correlation scores over the number of genes expressed within each subcluster. We performed Spearman rank correlations and show the lines of best fit in red.

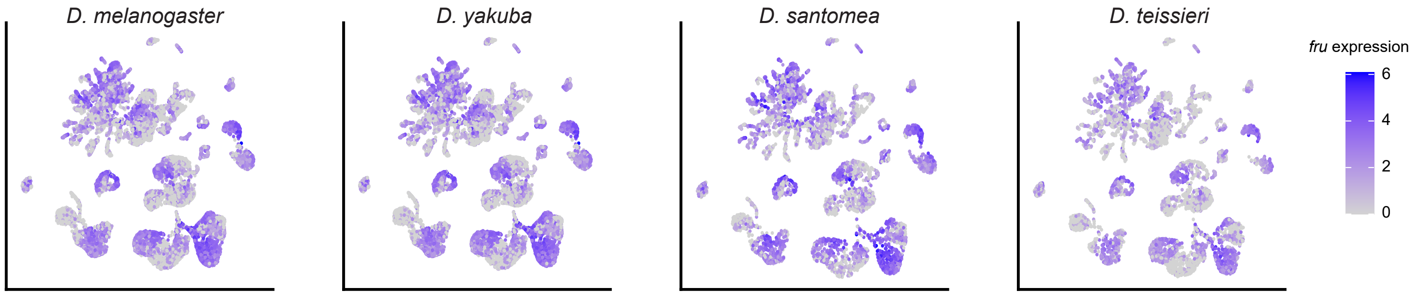

**Supplemental Figure 6**

Gene expression of *fru* across species in all *dsx*+ neurons.

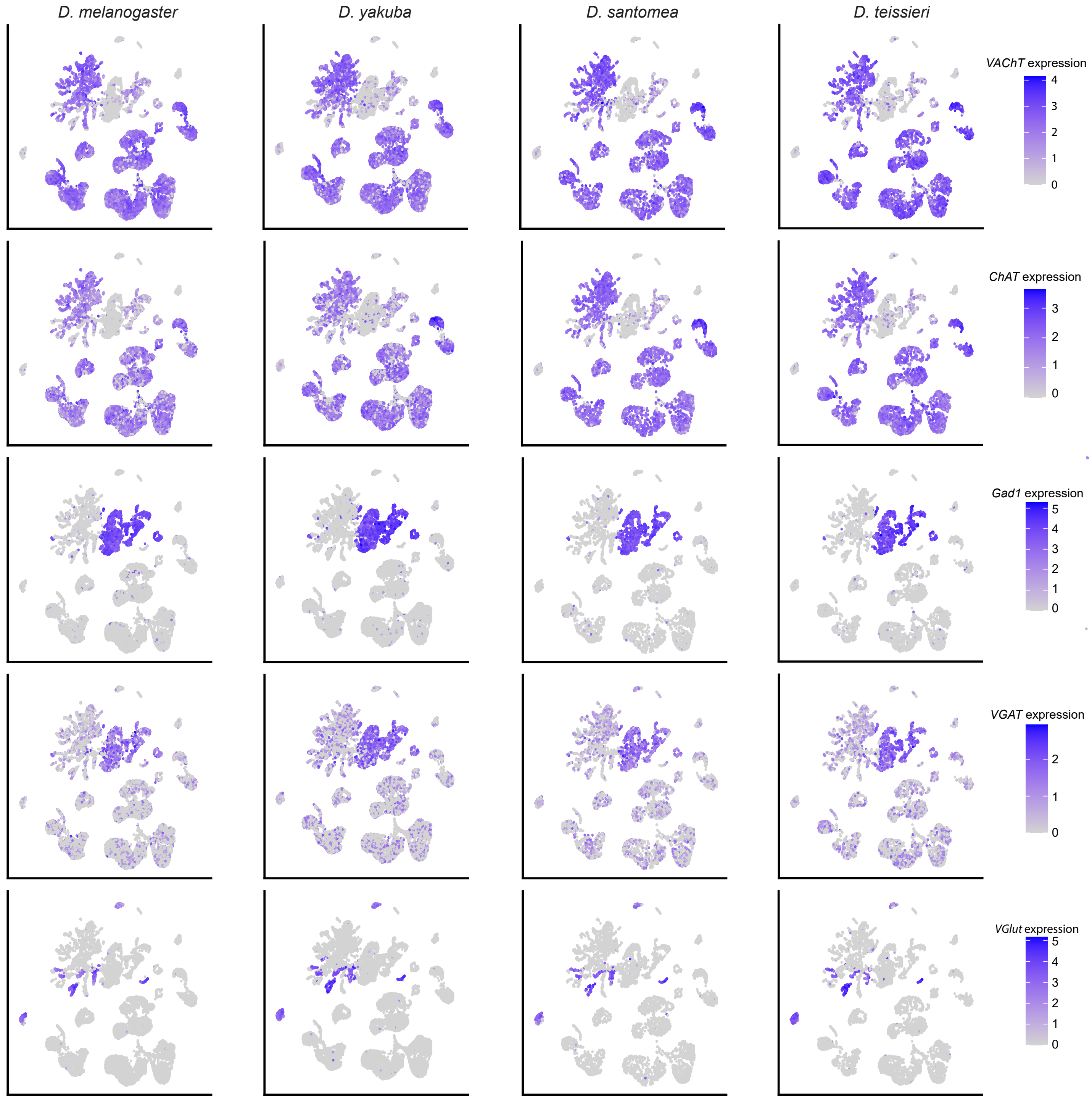

**Supplemental Figure 7**

Gene expression of *VAChT*, *ChAT*, *Gad1*, *VGAT*, and *VGlut* across species in all *dsx*+ neurons.

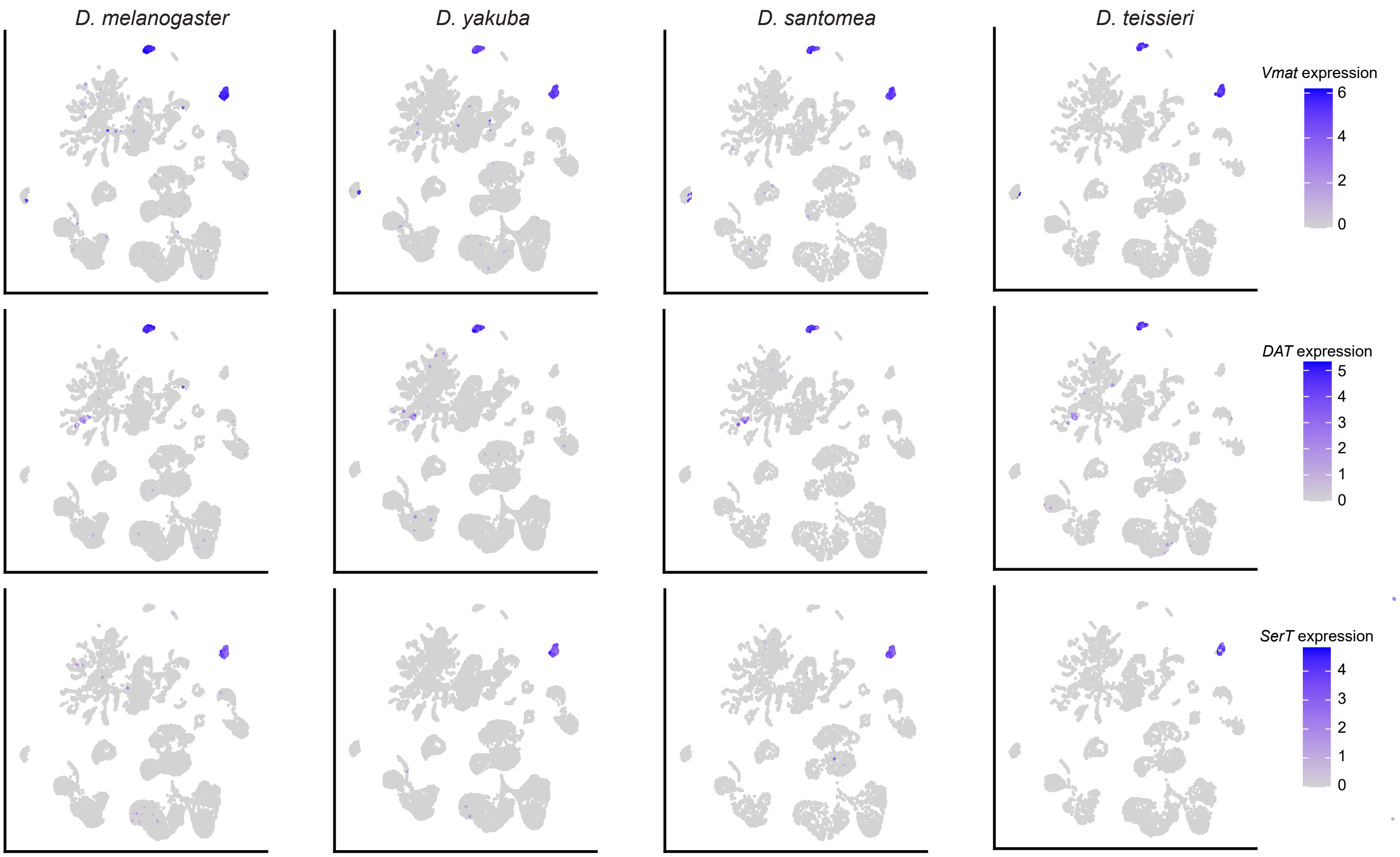

**Supplemental Figure 8**

Gene expression of *Vmat*, *DAT*, and *SerT* across species in all *dsx*+ neurons.

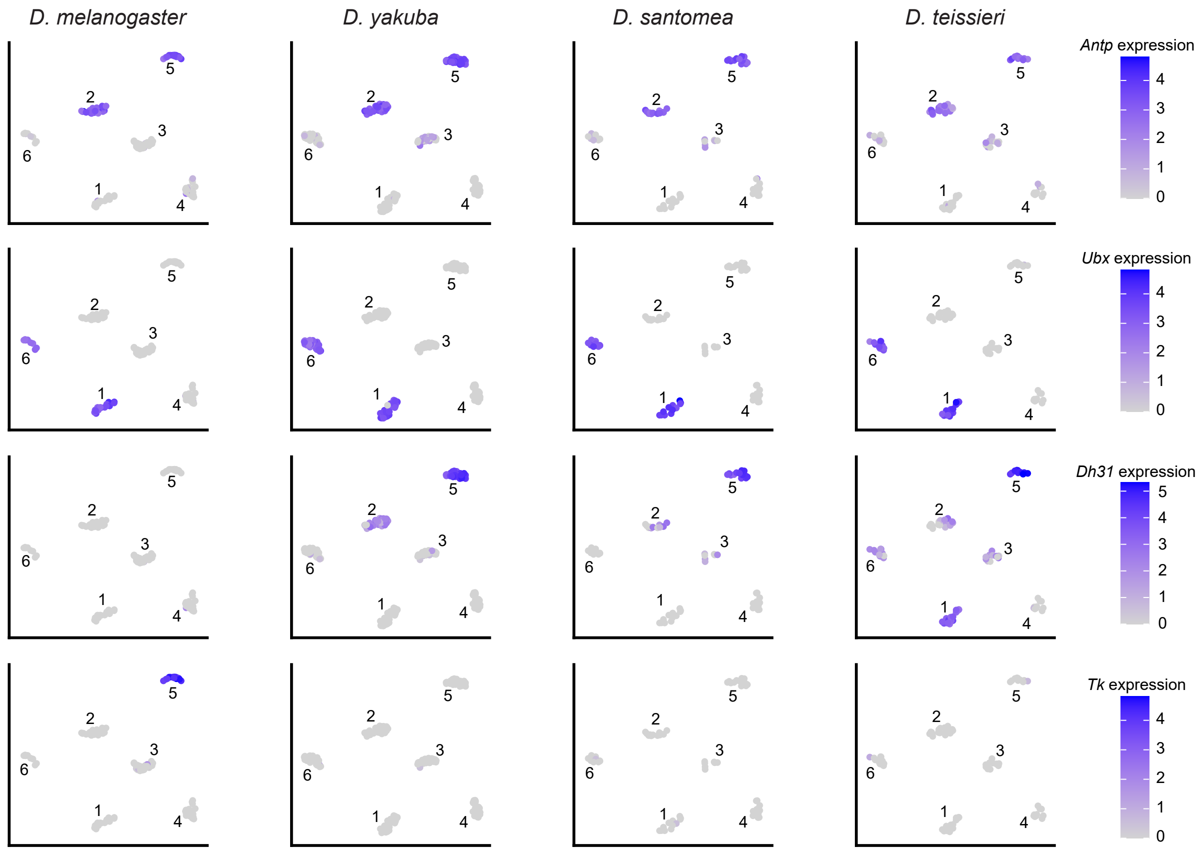

**Supplemental Figure 9**

Gene expression in TN2 of *Antp*, *Ubx*, *Dh31,* and *Tk* across species in TN2.

| **Supplemental Table 1. Summary of sequencing and filtering information.** | | | | | | |
|---|---|---|---|---|---|---|
| Species | Sex/ Replicate | % Seq saturation | % reads mapped | Reads per cell | # of cells | # of cells after filtering |
| *D. melanogaster* | M1 | 67.8 | 92.9 | 39,703 | 5,223 | 4,678 |
| *D. melanogaster* | M2 | 72.7 | 94.5 | 40,451 | 5,795 | 5,470 |
| *D. melanogaster* | M3 | 65.6 | 93.0 | 31,005 | 7,468 | 7,020 |
| *D. yakuba* | M1 | 61.1 | 90.3 | 30,390 | 5,882 | 5,193 |
| *D. yakuba* | M2 | 75.7 | 94.4 | 36,351 | 6,526 | 6,180 |
| *D. yakuba* | M3 | 84.4 | 95.3 | 35,501 | 6,996 | 6,694 |
| *D. santomea* | M1 | 81.6 | 93.2 | 48,419 | 5,746 | 5,343 |
| *D. teissieri* | M1 | 75.4 | 94.3 | 38,148 | 6,722 | 6,344 |
| *D. melanogaster* | F1 | 89.6 | 95.3 | 54,862 | 4,670 | 3,802 |

| **Supplemental Table 2. Hemilineage annotations.** | | |
|---|---|---|
| **Hemilineage** | **ANm subcluster(s)** | **Marker Genes expressed** |
| 0A | 42 | *en, inv, Gad1* |
| 1A | 26, 31, 41 | *Dr, VACht* |
| 1B/6A | 6, 12 | *Gad1, sens-2, vg* |
| 5B | 1, 2, 16 | *toy, vg* |
| 6B | 7 | *sens-2, vg* |
| 7B | 39 | *mab-21, unc-4* |
| 8B | 29, 45 | *C15, Lim3, VAChT* |
| 9A | 32 | *Dr, Gad1* |
| 9B | 44 | *Lim3, tup, VGlut* |
| 12A | 5 | *TfAP-2, unc-4* |
| 13A/19A | 3, 9, 10 | *Dbx, Fer2, Gad1* |
| 14A | 30 | *Dr, toy, VGlut* |
| 17A | 25 | *unc-4, Hmx* |
| 20A/22A | 27 | *B-H1, B-H2, VACht* |
| 21A | 28, 38 | *Dr, ey, VGlut* |
| 23B | 35 | *otp, slou, unc-4, VACht* |
| Unannotated | 4, 8, 11, 13, 14, 15, 17, 18, 19, 20, 21, 22, 23, 24, 33, 34, 36, 37, 40, 43, 46, 47, 48, 49, 50 | |

**Supplemental Table 3.** ***dsx*+ cell count.**

| Cluster | Species | Males Average ± SD | n | Females Average ± SD | n |
|---|---|---|---|---|---|
| aDN | *D. melanogaster* | 2.00 ± 0.00 | 9 | 2.00 ± 0.00 | 17 |
| | *D. teissieri* | 1.80 ± 0.52 | 20 | 2.00 ± 0.00 | 14 |
| | *D. yakuba* | 2.00 ± 0.00 | 9 | 2.00 ± 0.00 | 14 |
| | *D. santomea* | 2.00 ± 0.00 | 8 | 1.80 ± 0.52 | 20 |
| pC1 | *D. melanogaster* | 69.63 ± 1.92 | 8 | 9.35 ± 1.27 | 17 |
| | *D. teissieri* | 58.04 ± 5.39 | 24 | 4.93 ± 0.27 | 14 |
| | *D. yakuba* | 57.00 ± 2.05 | 10 | 5.36 ± 1.34 | 14 |
| | *D. santomea* | 71.13 ± 4.79 | 8 | 5.15 ± 0.59 | 20 |
| pC2l | *D. melanogaster* | 47.67 ± 1.66 | 9 | 13.80 ± 2.35 | 10 |
| | *D. teissieri* | 44.91 ± 3.78 | 23 | 11.64 ± 0.74 | 14 |
| | *D. yakuba* | 39.63 ± 1.69 | 8 | 9.79 ± 1.63 | 14 |
| | *D. santomea* | 47.57 ± 1.13 | 7 | 10.90 ± 1.37 | 20 |
| pC2m | *D. melanogaster* | 41.44 ± 3.28 | 9 | 1.38 ± 0.52 | 9 |
| | *D. teissieri* | 34.61 ± 3.97 | 9 | 1.21 ± 0.43 | 14 |
| | *D. yakuba* | 31.44 ± 1.94 | 8 | 2.00 ± 0.00 | 14 |
| | *D. santomea* | 31.63 ± 2.00 | 23 | 1.85 ± 0.49 | 20 |
| pCd-1 | *D. melanogaster* | 12.60 ± 0.97 | 10 | 7.56 ± 1.26 | 16 |
| | *D. teissieri* | 12.71 ± 0.95 | 24 | 7.79 ± 0.80 | 14 |
| | *D. yakuba* | 9.70 ± 0.92 | 10 | 7.21 ± 0.89 | 14 |
| | *D. santomea* | 12.38 ± 1.57 | 8 | 9.15 ± 0.88 | 20 |
| pCd-2 | *D. melanogaster* | 3.10 ± 0.32 | 10 | 3.00 ± 0.00 | 17 |
| | *D. teissieri* | 2.92 ± 0.28 | 24 | 3.00 ± 0.00 | 14 |
| | *D. yakuba* | 2.90 ± 0.32 | 10 | 2.93 ± 0.27 | 14 |
| | *D. santomea* | 2.86 ± 0.38 | 7 | 2.85 ± 0.75 | 20 |
| pLN | *D. melanogaster* | 1.00 ± 0.00 | 10 | 0 | 17 |
| | *D. teissieri* | 0.92 ± 0.28 | 24 | 0 | 14 |
| | *D. yakuba* | 1.00 ± 0.00 | 10 | 0 | 14 |
| | *D. santomea* | 1.00 ± 0.00 | 8 | 0 | 20 |
| pMN1 | *D. melanogaster* | 1.00 ± 0.00 | 10 | 1.00 ± 0.00 | 17 |
| | *D. teissieri* | 0.96 ± 0.20 | 24 | 1.00 ± 0.00 | 14 |
| | *D. yakuba* | 0.90 ± 0.32 | 10 | 1.00 ± 0.00 | 14 |
| | *D. santomea* | 1.00 ± 0.00 | 8 | 1.00 ± 0.00 | 20 |
| pMN2 | *D. melanogaster* | 0 | 10 | 1.00 ± 0.00 | 17 |
| | *D. teissieri* | 0 | 24 | 1.00 ± 0.00 | 14 |
| | *D. yakuba* | 0 | 10 | 1.00 ± 0.00 | 14 |
| | *D. santomea* | 0 | 8 | 1.00 ± 0.00 | 20 |
| pMN3 | *D. melanogaster* | 1.00 ± 0.00 | 10 | 0 | 17 |
| | *D. teissieri* | 1.00 ± 0.00 | 24 | 0 | 14 |
| | *D. yakuba* | 1.00 ± 0.00 | 10 | 0 | 14 |
| | *D. santomea* | 1.13 ± 0.35 | 8 | 0 | 20 |
| SN | *D. melanogaster* | 1.00 ± 0.00 | 9 | 0 | 17 |

| | | | | | |
|---|---|---|---|---|---|
| | *D. teissieri* | 0.74 ± 0.45 | 23 | 0 | 14 |
| | *D. yakuba* | 1.00 ± 0.00 | 10 | 0 | 14 |
| | *D. santomea* | 1.00 ± 0.00 | 6 | 0 | 20 |
| TN1* | *D. melanogaster* | 29.51 ± 1.80 | 49 | 0 | 17 |
| | *D. teissieri* | 24.64 ± 1.15 | 14 | 0 | 14 |
| | *D. yakuba* | 19.20 ± 1.86 | 60 | 0 | 14 |
| | *D. santomea* | 21.45 ± 0.95 | 20 | 0 | 20 |
| TN2 (total) | *D. melanogaster* | 7.45 ± 0.93 | 15 | 0 | 17 |
| | *D. teissieri* | 6.50 ± 0.71 | 2 | 0 | 14 |
| | *D. yakuba* | 7.78 ± 0.44 | 9 | 0 | 14 |
| | *D. santomea* | 8.00 ± 0.00 | 6 | 0 | 20 |
| TN2 (Pr only) | *D. melanogaster* | 3.92 ± 0.29 | 13 | 0 | 17 |
| | *D. teissieri* | 2.50 ± 0.58 | 4 | 0 | 14 |
| | *D. yakuba* | 4.00 ± 0.00 | 10 | 0 | 14 |
| | *D. santomea* | 4.00 ± 0.00 | 14 | 0 | 20 |
| TN2 (Ms only) | *D. melanogaster* | 2.55 ± 0.69 | 11 | 0 | 17 |
| | *D. teissieri* | 3.00 ± 0.00 | 4 | 0 | 14 |
| | *D. yakuba* | 2.80 ± 0.42 | 10 | 0 | 14 |
| | *D. santomea* | 3.00 ± 0.00 | 14 | 0 | 20 |
| TN2 (Mt only) | *D. melanogaster* | 1.00 ± 0.00 | 12 | 0 | 17 |
| | *D. teissieri* | 1.00 ± 0.00 | 2 | 0 | 14 |
| | *D. yakuba* | 1.00 ± 0.00 | 9 | 0 | 14 |
| | *D. santomea* | 1.00 ± 0.00 | 10 | 0 | 20 |

All counts taken from one side of the brain/ventral nerve cord

*data from Ye, Walsh et al., 2024

**Supplemental Table 4. Subcluster matches between the male/female and four-species datasets.**

| Male/Female Cluster | Four Species Cluster | Percent Match |
|---|---|---|
| fm_aDN_1 | aDN_1 | 73.8 |
| | aDN_2 | 26.2 |
| fm_pC1_1 | pC1_1 | 99.2 |
| fm_pC1_2 | pC1_3 | 99.2 |
| fm_pC1_3 | pC1_2 | 99.4 |
| fm_pC1_4 | pC1_4 | 98.2 |
| fm_pC1_5 | pC1_5 | 98.1 |
| fm_pC1_6 | pC1_6 | 94.3 |
| fm_pC2l_1 | pC2l_1 | 99.6 |
| fm_pC2l_2 | pC2l_2 | 57.5 |
| | pC2l_5 | 30.9 |
| | pC2l_6 | 10.3 |
| fm_pC2l_3 | pC2l_3 | 99.1 |
| fm_pC2l_4 | pC2l_1 | 99.0 |
| fm_pC2l_5 | pC2l_4 | 97.0 |
| fm_pC2l_6 | pC2l_2 | 98.5 |
| fm_pC2m_1 | pC2m_1 | 96.1 |
| fm_pC2m_2 | pC2m_2 | 99.4 |
| fm_pC2m_3 | pC2m_3 | 99.0 |
| fm_pC2m_4 | pC2m_4 | 98.7 |
| fm_pCd1_1 | pCd1_1 | 57.4 |
| | pCd1_3 | 41.6 |
| fm_pCd1_2 | pCd1_2 | 99.3 |
| fm_pCd1_3 | pCd1_1 | 100 |
| fm_pCd2_1 | pCd2_2 | 100 |
| fm_pCd2_2 | pCd2_1 | 100 |
| fm_pCd2_3 | pCd2_3 | 93.3 |
| fm_TN1_1 | TN1_2 | 97.4 |
| fm_TN1_2 | TN1_1 | 98.8 |
| fm_TN1_3 | TN1_3 | 96.0 |
| fm_TN1_4 | TN1_1 | 53.9 |
| | TN1_4 | 45.1 |
| fm_TN2_1 | TN2_3 | 100 |
| fm_TN2_2 | TN2_4 | 100 |
| fm_TN2_3 | TN2_1 | 73.3 |
| | TN2_2 | 26.7 |
| fm_TN2_4 | TN2_5 | 100 |
| fm_ANm_1 | ANm_1 | 88.2 |
| fm_ANm_2 | ANm_17 | 39.2 |
| | ANm_43 | 14.4 |

| | | |
|---|---|---|
| | ANm_21 | 12.4 |
| fm_ANm_3 | ANm_3 | 85.1 |
| | ANm_9 | 14.9 |
| fm_ANm_4 | ANm_6 | 100 |
| fm_ANm_5 | ANm_24 | 62.3 |
| | ANm_22 | 29.0 |
| fm_ANm_6 | ANm_5 | 98.9 |
| fm_ANm_7 | ANm_2 | 98.8 |
| fm_ANm_8 | ANm_10 | 100 |
| fm_ANm_9 | ANm_23 | 79.3 |
| | ANm_8 | 19.5 |
| fm_ANm_10 | ANm_4 | 100 |
| fm_ANm_11 | ANm_14 | 100 |
| fm_ANm_12 | ANm_11 | 85.1 |
| | ANm_25 | 14.9 |
| fm_ANm_13 | ANm_13 | 97.3 |
| fm_ANm_14 | ANm_9 | 100 |
| fm_ANm_15 | ANm_7 | 99.1 |
| fm_ANm_16 | ANm_16 | 89.1 |
| | ANm_1 | 10.0 |
| fm_ANm_17 | ANm_15 | 100 |
| fm_ANm_18 | ANm_1 | 92.1 |
| fm_ANm_19 | ANm_5 | 100 |
| fm_ANm_20 | ANm_8 | 100 |
| fm_ANm_21 | ANm_20 | 100 |
| fm_ANm_22 | ANm_12 | 98.7 |
| fm_ANm_23 | ANm_29 | 92.9 |
| fm_ANm_24 | ANm_18 | 95.7 |
| fm_ANm_25 | ANm_26 | 100 |
| fm_ANm_26 | ANm_17 | 64.3 |
| | TN2_3 | 14.3 |
| fm_ANm_27 | ANm_12 | 100 |
| fm_ANm_28 | ANm_28 | 100 |
| fm_ANm_29 | ANm_21 | 100 |
| fm_ANm_30 | ANm_36 | 90.1 |
| fm_ANm_31 | ANm_30 | 100 |
| fm_ANm_32 | ANm_35 | 91.1 |
| fm_ANm_33 | ANm_31 | 100 |
| fm_ANm_34 | ANm_41 | 92.6 |
| fm_ANm_35 | ANm_22 | 100 |
| fm_ANm_36 | ANm_40 | 71.1 |
| | ANm_47 | 28.9 |
| fm_ANm_37 | ANm_38 | 97.2 |
| fm_ANm_38 | ANm_46 | 100 |

| | | |
|---|---|---|
| fm_ANm_39 | ANm_42 | 97.6 |
| fm_ANm_40 | ANm_32 | 79.5 |
| | ANm_50 | 20.5 |
| fm_ANm_41 | Female specific | NA |
| fm_ANm_42 | ANm_34 | 97.7 |
| fm_ANm_43 | ANm_39 | 100 |
| fm_ANm_44 | ANm_37 | 100 |
| fm_ANm_45 | ANm_33 | 100 |
| fm_ANm_46 | ANm_44 | 96.2 |
| fm_ANm_47 | ANm_27 | 93.3 |
| fm_ANm_48 | ANm_4 | 96.6 |
| fm_ANm_49 | Female specific | NA |
| fm_ANm_50 | ANm_3 | 100 |

**Supplemental Table 5. List of genotypes, reagents, and software used in this study.**

| REAGENT or RESOURCE | SOURCE | IDENTIFIER |
|---|---|---|
| **Chemicals, peptides, and recombinant proteins** | | |
| Liberase DH | Thermo Fisher Scientific | Cat# 50-100-3341 |
| Schneider's *Drosophila* medium | Thermo Fisher Scientific | Cat# 21-720-024 |
| Bovine Serum Albumin | Thermo Fisher Scientific | Cat# B14 |
| **Critical Commercial Assays** | | |
| **Deposited Data** | | |
| scRNA-seq data | GEO Accession | N/A |
| **Experimental Models: Organisms/Strains** | | |
| *D. melanogaster*, ; pJFRC29-10XUAS-IVS-myr::GFP-p10 (attP40)/+; pJFRC105-10XUAS-IVS-nls::tdTomato (VK00040)/+ | Gerry Rubin's lab | N/A |
| *D. yakuba*, w; ; pJFRC105-10XUAS-nls::tdTomato (2283) | Ye et al., 2024 | N/A |
| *D. santomea*, w; ; pJFRC105-10XUAS-nls::tdTomato (2253) | This study | N/A |
| *D. teissieri*, pBac{10XUAS-nls::tdTomato, 3XP3::YFP} | This study | N/A |
| *D. melanogaster*, ; ; dsx-GAL4/UAS-myrGFP | Ye et al., 2024 | N/A |
| *D. yakuba*, ; ; dsx-GAL4/UAS-myrGFP | Ye et al., 2024 | N/A |
| *D. santomea*, ; UAS-myrGFP/+; dsx-GAL4/+ | Ye et al., 2024 | N/A |
| *D. teissieri*, ; UAS-myrGFP/+; dsx-GAL4/+ | Ye et al., 2024 | N/A |
| *D. melanogaster*, w+;;*tey*-vp16#ex1/TM6B | This study; see Chen et al., 2023 for method details | N/A |
| *D. melanogaster*, w; *optix*-DBD; | Simon et al., 2024 | N/A |
| *D. melanogaster*, ; ; *TfAP-2*-AD/TM6B | Ye et al., 2024 | N/A |
| *D. melanogaster*, *shaven*-AD/ln(4)d | Soffers et al., 2025 | N/A |
| *D. melanogaster*, *unc-4*-AD/FM7; ; | Lacin et al., 2020 | N/A |
| *D. melanogaster*, ; sp/CyO; Gad1-DBD/TM6B | Lacin et al., 2020 | N/A |
| *D. melanogaster*, ; ; *ara*-GAL4DBD#ex1/TM6B | This study; see Chen et al., 2023 for method details | N/A |
| *D. melanogaster*, w; ; *dsx*-DBD(w+)/TM6B | Shirangi et al., 2016 | N/A |
| *D. melanogaster*, ; if/CyO; *dsx*-vp16 #M3/TM6B | This study; see Chen et al., 2023 for method details | N/A |
| **Oligonucleotides** | | |
| mel-*dsx*-B1 HCR probe set | Ye et al., 2024 | N/A |

| | | |
|---|---|---|
| yak-*dsx*-B1 HCR probe set | Ye et al., 2024 | N/A |
| mel-*Dh31*-B3 HCR probe set | Molecular Instruments | N/A |
| mel-T*k*-B2 HCR probe set | Molecular Instruments | N/A |
| B1 AF647 HCR Amplifier | Molecular Instruments | N/A |
| B2 AF488 HCR Amplifier | Molecular Instruments | N/A |
| B3 AF546 HCR Amplifier | Molecular Instruments | N/A |
| **Software and Algorithms** | | |
| ZEN Digital Imaging for Light Microscopy | Zeiss | RRID: SCR_013672 |
| Leica Application Suite X | Leica | RRID: SCR_013673 |
| Adobe Illustrator | Adobe Systems | RRID: SCR_010279 |
| VVD Viewer | https://github.com/takashi310/VVD_Viewer/blob/master/README.md | N/A |
| Fiji | NIH. https://imagej.net/fiji | RRID: SCR_002285 |
| RStudio | R Core Team https://posit.co/ | RRID: SCR_000432 |
| 10x Genomics Cell Ranger | Zheng et al.[71] | RRID: SCR_017344 |
